# Comparing the utility of *in vivo* transposon mutagenesis approaches in yeast species to infer gene essentiality

**DOI:** 10.1101/732552

**Authors:** Anton Levitan, Andrew N. Gale, Emma K. Dallon, Darby W. Kozan, Kyle W. Cunningham, Roded Sharan, Judith Berman

**Affiliations:** School of Molecular Microbiology and Biotechnology, Faculty of Life Sciences, Tel Aviv University, Tel Aviv, Israel; Edmond J. Safra Center for Bioinformatics, Tel Aviv University, Tel Aviv, Israel; Department of Biology, Johns Hopkins University, Baltimore, MD, USA; Blavatnik School of Computer Science, Tel Aviv University, Tel Aviv, Israel

**Keywords:** Genomics, Yeasts, Bioinformatics, Machine Learning, High-throughput

## Abstract

*In vivo* transposon mutagenesis, coupled with deep sequencing, enables large-scale genome-wide mutant screens for genes essential in different growth conditions. We analyzed six large-scale studies performed on haploid strains of three yeast species (*Saccharomyces cerevisiae, Schizosaccaromyces pombe*, and *Candida albicans*), each mutagenized with two of three different heterologous transposons (*AcDs, Hermes*, and *PiggyBac*). Using a machine-learning approach, we evaluated the ability of the data to predict gene essentiality. Important data features included sufficient numbers and distribution of independent insertion events. All transposons showed some bias in insertion site preference because of jackpot events, and preferences for specific insertion sequences and short-distance vs long-distance insertions. For *PiggyBac*, a stringent target sequence limited the ability to predict essentiality in genes with few or no target sequences. The machine learning approach also robustly predicted gene function in less well-studied species by leveraging cross-species orthologs. Finally, comparisons of isogenic diploid versus haploid *S. cerevisiae* isolates identified several genes that are haplo-insufficient, while most essential genes, as expected, were recessive. We provide recommendations for the choice of transposons and the inference of gene essentiality in genome-wide studies of eukaryotic haploid microbes such as yeasts, including species that have been less amenable to classical genetic studies.

## INTRODUCTION

Work with model yeasts such as *Saccharomyces cerevisiae* and *Schizosaccharomyces pombe* has pioneered the combination of genotype/phenotype comparisons at a genomic scale. Work on these yeasts, which have had genome sequences available for more than 20 years (Goffeau *et al*., 1996; Wood *et al*., 2002), along with facile gene replacement protocols, has relied heavily on comprehensive collections of deletion mutants (Giaever *et al*., 2002; Kim *et al*., 2010) for high-throughput dissection of specific genotypes, as well as for genetic interactions with drugs (reviewed in (Lehár *et al*., 2008) and gene-gene interactions through systematic analysis of double and triple mutant analysis (e.g., (Reguly *et al*., 2006); (Kuzmin *et al*., 2018). While genome wide deletion collections are very useful for systematic studies, some individual deletion mutants acquire additional mutations or copy number variations in genes, genome regions, chromosomes or mitochondrial DNA, at least some of which improve the fitness of the original mutations (Hughes *et al*., 2000; Ben-Shitrit *et al*., 2012; Yona *et al*., 2012; Puddu *et al*., 2019).

In animals and plants that are less amenable to such directed molecular manipulations, the use of heterologous transposons *in vivo* has facilitated genetic analysis, within the limitations imposed by the transposon excision/insertion process (Munoz-Lopez and Garcia-Perez, 2010; Kawakami, Largaespada and Ivics, 2017). With the advent of deep sequencing, such studies have become more facile and have been performed in the two model yeasts as well (Gangadharan *et al*., 2010; Guo *et al*., 2013; Michel *et al*., 2017). *In vivo* transposon mutagenesis generally involves the introduction of a heterologous DNA transposon, along with the genes (e.g., the relevant transposase) required to induce its active transposition into a clonal isolate of a species of interest. Upon induction, the transposase excises the transposon from its original location (excision site) and inserts it into a single new position in the genome. Because the frequency of transposon excision and reinsertion is quite low, each cell harbors, at most, a single transposition mutation. The transposon is usually engineered for facile selection of excision and/or reinsertion events, allowing detection and enrichment of these rare events.

*In vivo* transposon insertion provides several advantages, as it rapidly yields large numbers of mutants in a single step and easily can be performed in parallel strains with different mutations or genetic backgrounds. Because it does not require much prior knowledge, it also can be performed in non-model species, where each experiment is likely to provide much new information. The only transformation steps required are those used to engineer the starting strain. This bypasses the problem of low transformation efficiency in many species. It also avoids the unintended genome alterations (e.g., aneuploidies) that often accompany DNA transformation (Bouchonville *et al*., 2009). The sites of transposon insertion throughout the genome can be identified *en masse* using very large collections of independent insertion event clones, coupled with deep sequencing of the DNA immediately adjacent to the new transposon locus (Wu *et al*., 2007; Rad *et al*., 2010; Yusa *et al*., 2011). For example, a high throughput genotype/phenotype analysis of 30 bacterial species grown under >170 different nutrient and stress conditions recently assigned functions to thousands of genes, including ∼300–600 genes per bacterium that are essential for viability (Price *et al*., 2018).

Three different transposon systems have been used for *in vivo* mutagenesis in yeasts: *AcDs* from *Zea mays, Hermes* from *Musca domestica*, and *PiggyBac* from *Trichoplusia ni. AcDs* has been used primarily in plant species, but was also engineered for increased efficiency in the model yeast *S. cerevisiae* (Lazarow *et al*., 2012; Michel *et al*., 2017) and, later, in *C. albicans (Mielich et al*., *2018). AcDs* does not display a strong insertion sequence preference, although it has a higher frequency of insertions into intergenic regions than into coding regions and has a bias for reinsertion near the initial site of excision. *Hermes* has been used for mutagenesis in *S. pombe* and *S. cerevisiae* (Park, Evertts and Levin, 2009; Gangadharan *et al*., 2010; Edskes *et al*., 2018), it prefers to insert at genomic positions with the target sequence TnnnnA. *PiggyBac* (PB) has been used in mammalian systems such as rat and mouse (Rad *et al*., 2010; Yusa *et al*., 2011; Zhao *et al*., 2016), and also in *S. pombe (Li et al*., *2011). PB* has a strong preference for insertion at TTAA sequences, which are generally more frequent in AT-rich intergenic regions than within coding sequences.

Transposon insertion within an ORF is generally assumed to cause a loss-of-function mutation. Identifying the phenotypes associated with loss-of-function mutations in specific genes allows the prediction of genetic functions. Cells in which the transposon inserted into a gene essential for viability will fail to grow and thus be lost from the population. By contrast, cells with mutations in non-essential genes are expected to be well-represented in the cell population. Insertion of a transposon carrying a strong promoter into an ORF could activate expression inappropriately; this can be useful for the study of gain-of-function mutations.

*In vivo* transposon mutagenesis studies of yeasts include analysis of *S. cerevisiae* with *Hermes* (this study and (Park, Evertts and Levin, 2009; Gangadharan *et al*., 2010; Edskes *et al*., 2018)) or with the mini-Ds derivative of the *AcDs* system (Michel *et al*., 2017), in *S. pombe* with *Hermes* (Guo *et al*., 2013) and *PB* (Li *et al*., 2011), and in *C. albicans* with *AcDs (Segal et al*., *2018)* and with *PB* (*Gao et al*., *2018*) (Fig. 1a). In earlier work, we applied a machine learning (ML) approach to infer gene essentiality from the *C. albicans AcDs* data. Here, we modified that ML approach to predict the likelihood of essentiality for the complete set of predicted open reading frames for these six *in vivo* transposon datasets. We compared the strengths and challenges of the different transposons in each species, looking for insights into the number of insertion events required for accurate predictions, the distribution of mutations, and the degree to which different transposons, with different sequence dependencies, provided similar or different conclusions. The goals of this study were to provide metrics that could assist in determining the advantages and disadvantages of different transposon systems, so as to optimize the data produced in a given *in vivo* transposon system, and to suggest approaches for generating whole genome data in understudied yeast species.

## MATERIALS AND METHODS

### Data Acquisition

#### 1. Experimental

*Sc Hermes* data was obtained as follows: haploid and diploid strains of *S. cerevisiae* were transformed with plasmid pSG36 (Gangadharan *et al*., 2010). A single colony was suspended in 100 mL synthetic complete (SC) medium lacking uracil and containing 2% galactose, divided into twenty 16 x 150 mm glass culture tubes, and shaken for 3 days at 30°C. This protocol yielded ∼5 x 10^6 cells bearing transposon insertions per mL (∼3% of all cells). To enrich for cells bearing transposon insertions, the twenty cultures were pooled and centrifuged, and the cell pellet was resuspended in 600 mL SC medium containing 2% glucose, 0.1 mg/mL nourseothricin, and 1 mg/mL 5-fluoroorotic acid, and then shaken overnight at 30°C. The cells were pelleted, resuspended in 600 mL of the same medium, and cultured as before. Finally, 60 mL of these enriched cells were pelleted, resuspended in 600 mL of the same medium, and cultured as before. These highly enriched cells were pelleted, resuspended in 15% glycerol, and frozen in aliquots at-80°C. To extract genomic DNA, 100 mg of thawed cell pellets were washed three times in 1 mL deionized water and extracted using Quick-DNA Fungal/Bacterial Miniprep kit (Zymo Research). A total of 2.4 µg of purified gDNA was fragmented by sonication in four separate tubes using a Diagenode Picoruptor. The fragmented DNA was then end repaired, ligated to splinkerette adapters, size selected with AMPure xp beads, and PCR amplified in separate reactions using transposon-specific and adapter-specific primers, as detailed previously (Bronner *et al*., 2016).

Samples were then PCR amplified to attach Illumina P5 and P7 (indexed) adapters, purified with AMPure xp beads, mixed with phiX-174, and loaded into the MiSeq instrument (Illumina); 75 bp of each end was then sequenced using primers specific for Hermes right inverted repeat and P7. Detailed protocols and primer sequences are available upon request. De-multiplexed reads were mapped to the *S. cerevisiae* S288C reference genome using Bowtie2 (Langmead and Salzberg, 2012), and any mapped reads with a quality score < 20 or a mismatch at nucleotide +1 were removed. This process was repeated a total of 3 times in diploid strain BY4743, 2 times in haploid strain BY4741, and 1 time in haploid strain BY4742. The diploid and haploid datasets were combined prior to analyses. The *S*.*cerevisiae Hermes* data (mapped reads and counts) are available at http://genome-euro.ucsc.edu/s/CunninghamLab/Hermes%20Vs%20AcDs.

**Fig. 1.**
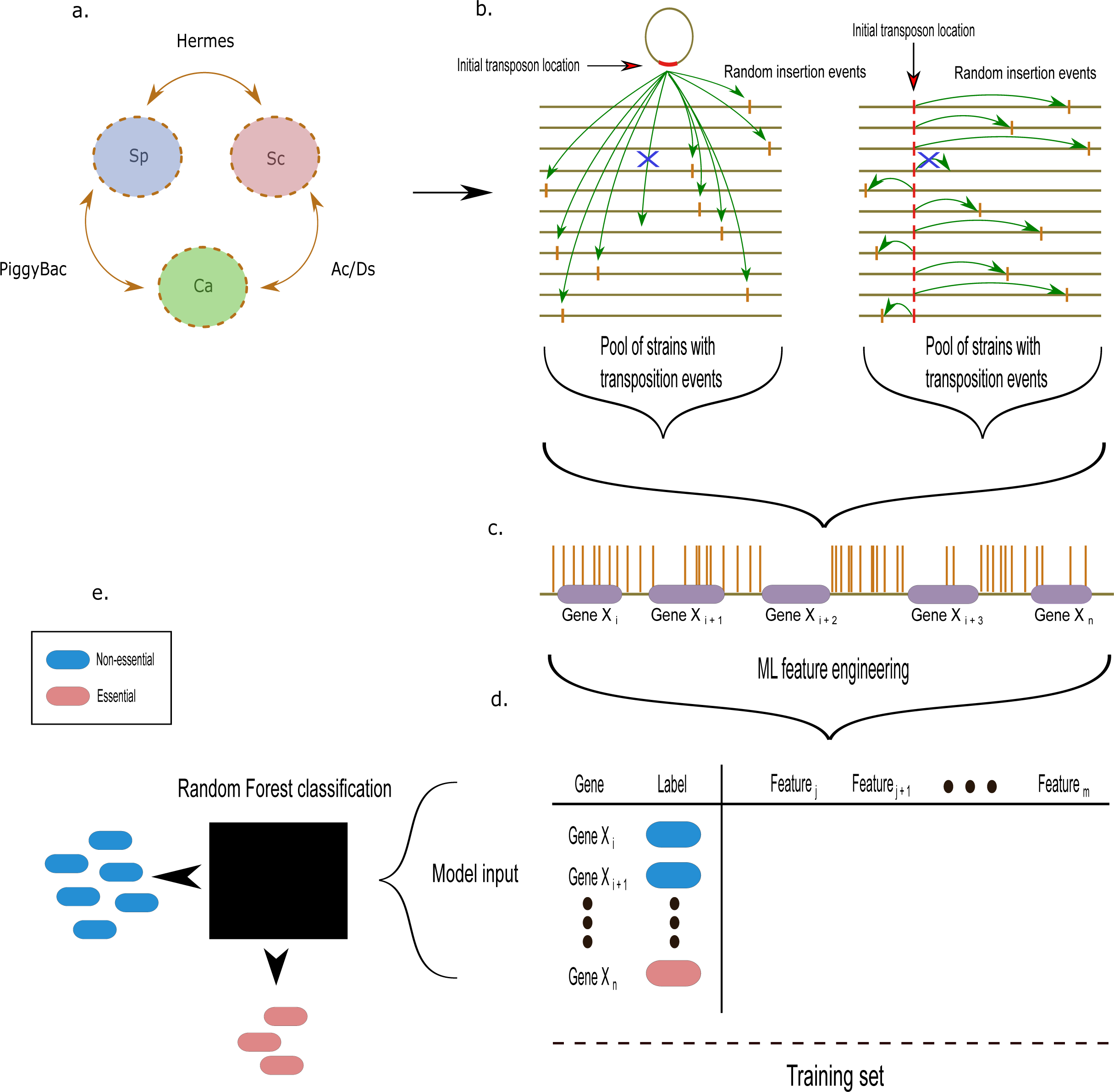
Overview of data acquisition and analysis: transposition events to gene essentiality. **a.** Three yeast species analyzed (Sp, *S. pombe*; Sc, *S. cerevisiae*; and Ca, *C. albicans*) by *in vivo* transposition in this study and two out of the three transposons (PB, *PiggyBac*; *AcDs* and *Hermes*) were to mutagenize each species. Note that each species was analyzed with two different transposon systems. **b.** Comparison of genome insertion sites for transposition events that initiate from an extrachromosomal plasmid (left, red region of plasmid circle) or a specific chromosomal locus (right, red bar on a given chromosome). Horizontal lines represent multiple copies of the same genome, each of which underwent a single insertion event (green arrow) per genome. While transposition is generally random, a bias for loci in close proximity to the initial transposon insertion site demands normalization of the final data. **c.** Mapped Tnseq analysis of the pool of transposition events yields the chromosomal insertion sites (brown vertical lines) in the reference chromosomes relative to the ORFs (purple regions). A close up of a small region of a single chromosome (olive horizontal line) including 5 ORFs is illustrated. **d.** A training set is constructed using known or inferred labels (non-essential, blue; essential, red) together with extracted features calculated from the data and its position relative to ORFs. Features are defined in Table 1. **e.** The training set features, as well as features for all ORFs are used as input for Random Forest classification (black rectangle); output is a prediction of essentiality (red or blue as in d), for which an optimal threshold is determined and applied to designate all genes in one of the two categories.

#### 2. Publicly available databases

The rest of the datasets analyzed here were obtained from previously published studies. *ScAcDs*, from which both WT1 & WT2 were combined for the analysis, was published by (Michel *et al*., 2017). The data was downloaded from https://www.ebi.ac.uk/arrayexpress/experiments/E-MTAB-4885/samples/. *SpPB* was published by (Li *et al*., 2011) and the data was obtained from the SRA database (Leinonen *et al*., 2011): SRR089408. *Sc Hermes* was also published by (Park, Evertts and Levin, 2009; Gangadharan *et al*., 2010; Edskes *et al*., 2018), and this data was added to the analysis and is found in Table S4b. *Sp Hermes* was published by (Guo *et al*., 2013) and the data was obtained from the SRA database: SRR327340. *CaAcDs* was published by (*Segal et al*., *2018*) and the data was obtained from the SRA database, where the SRR7824843, SRR7824841, and SRR7824838 files were combined for the analysis. *CaPB* was published by (*Gao et al*., *2018*) and the data was obtained from the SRA database, where all following files were used for the analysis: DMSO (untreatment): SRR7704188, SRR7704193, SRR7704196; 5-FOA (untreatment: SRR7704189, SRR7704194, SRR7704200; No drug screen: SRR7704195). All SRR files were obtained using fastq-dump (*SRA database, fastq-dump software*, no date), without the technical reads and by splitting the forward and reverse reads into individual files, as follows: fastq-dump - -gzip --skip-technical --readids --dumpbase --split-files --clip <SRR*******>

**Table 1.** Features used in the Random Forest model for the inference of gene essentiality.

| Feature | Description |
| --- | --- |
| Insertions | Number of transposon insertions within the ORF |
| Reads | Number of reads associated with the transposon insertions within the ORF |
| Neighborhood Index (NI) | Number of transposon insertions within the ORF, normalized by the length of the ORF and the surrounding 10kbp |
| Freedom Index (FI) | Length of the largest insertion-free region in the ORF, divided by the ORF's length |
| Insertions 100 upstream | Number of transposon insertions within the upstream region of the ORF |
| Insertions per Target Seqs | Number of transposon insertions divided by the number of transposon target sequences within an ORF |
| Reads per Length | Number of transposon insertions divided by the length of the ORF |
| Insertions per Length | Number of reads associated with the transposon insertions divided by the length of the ORF |

### Data Processing

The fastq files, downloaded with fastq-dump, were processed further using cutadapt (Martin, 2011) to filter out reads not containing partial transposon sequences. Reads with transposon sequences were trimmed to remove the transposon sequences for alignment purposes, as follows: cutadapt -- cores=8 -m 2 -g <primer sequence> <input fastq filename> -o <output fastq filename> --discard-untrimmed --overlap <overlap length>. In the analysis of the *Sc Hermes* study, all the sequencing reads contained the transposon and the reads start at the first genomic base, thus requiring no filtering (similarly, for the *S. cerevisiae Hermes* data from (Edskes *et al*., 2018)). In the analysis of the *Sp PiggyBac*, we filtered the reads containing the transposons from the rest by identifying the ACGCAGACTATCTTTCTAGGG sequence, cutting it out, and aligning only the remaining part of the relevant reads. In the analysis of the *Ca AcDs*, we filtered the reads containing the transposons from the rest by identifying the GTATTTTACCGACCGTTACCGACC sequence, cutting it out, and aligning only the remaining part of the relevant reads (Segal *et al*., 2018). In the analysis of the *Ca PiggyBac*, we filtered the reads containing the transposons from the rest by identifying the TGCATGCGTCAATTTTACGCAGACTATCTTTCTA sequence, cutting it out, and aligning only the remaining part of the relevant reads, starting 3bp downstream. In the analysis of the *Sc AcDs* study, we used the published transposon insertion maps of WT1 and WT2. In the analysis of the *Sp Hermes* study, we used the published transposon insertion maps (Segal *et al*., 2018).

### Alignment of reads and mapping the transposon insertions

Bowtie2 (Langmead and Salzberg, 2012) indices were created for each organism and gffutils databases were created for each organism’s genetic features, using the latest versions of the reference genomes (fasta) and the genomic feature files (gff), which were downloaded from the respective official sources for the three organisms: *S. cerevisiae* from https://downloads.yeastgenome.org, *S. pombe* from ftp://ftp.pombase.org/pombe/ and *C. albicans* http://www.candidagenome.org/download/. Sequencing reads were aligned using Bowtie2 with the default settings. The resulting sam files were converted to bam using samtools (Li *et al*., 2009). bam files were sorted using samtools and indexed via pysam(*pysam: a Python library for SAM accessing files*, no date). Transposon insertions and their corresponding reads were mapped to the respective genomes and counted in each genomic feature. Transposon target sites were found in every genome using Biopython (Cock *et al*., 2009) and counted in each genetic feature.

### Gene essentiality classification

Table 1 summarizes the features for machine learning classification that were engineered from the mapped transposon insertions and reads and from the transposon target sequences in the genomes. Random Forest (Breiman, 2001) classification was performed, using Python’s scikit-learn library (Hao and Ho, 2019), with the default parameters, except the n_estimators parameter was increased to 200 and the random_state parameter was fixed at 0, for reproducibility purposes. The results were validated using a 5-fold cross-validation. Essentiality labels for the training set of each organism were obtained previously (Segal *et al*., 2018) and are provided in Table S1.

Thresholds for the essentiality predictions in each classification were chosen as follows: Two metrics were evaluated (Fig. 2): 1) Minimum of the Euclidean distance between (0, 1) and the receiver operating characteristic (ROC) curve. 2) Maximum of the vertical distance between the line describing a random choice (a straight line from (0, 0) to (1, 1)) and the ROC curve (Youden, 1950; Fluss, Faraggi and Reiser, 2005). The second method (Youden Index (Youden, 1950)) was chosen, and we verified that the first metric is reasonably close, to eliminate any possible artifacts. We predicted the essentiality of all the available genes for each organism based on their respective features, and used the aforementioned method to choose the threshold for each binary classification.

**Fig. 2.**
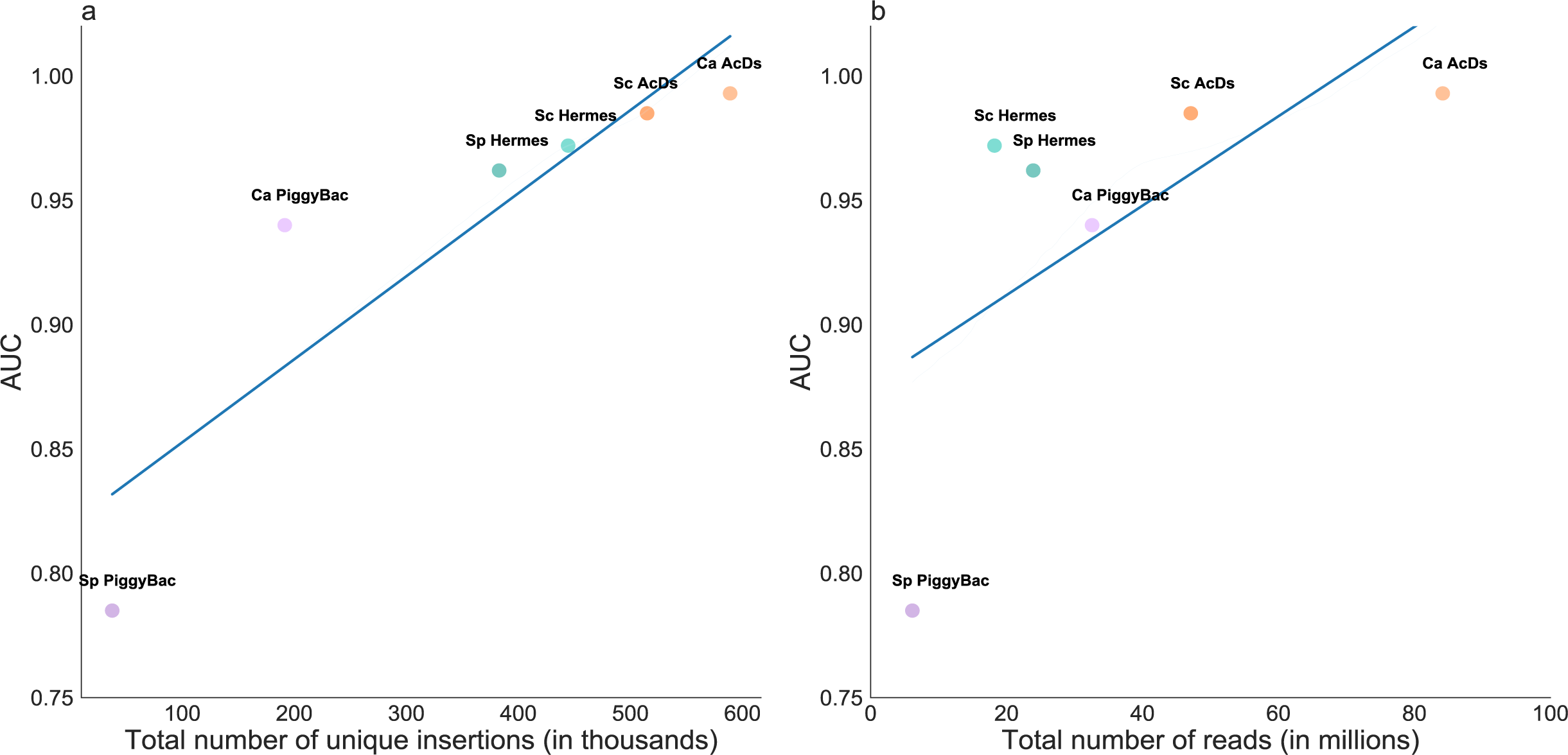
Threshold optimization. Two metrics were evaluated (Youden, 1950): 1) Minimum of the Euclidean distance between (0, 1) and the receiver operating characteristic (ROC) curve. 2) Maximum of the vertical distance between the line describing a random choice (a straight line from (0, 0) to (1, 1)) and the ROC curve (Youden Index).

### Manual curation of genes

First, we removed the genes that suffer technical limitations, which were primarily duplicated or repeated regions of the genome (Fig. 3) (Segal *et al*., 2018). Briefly, this is because when the aligner encounters more than one region for a possible successful alignment, it assigns a low score to this alignment and eliminates those sequences in the quality assurance phase. Thus, genes within duplicated regions appear to have few if any insertion sites (hits) and are falsely predicted to be essential by the classifier (Table S10). Second, we removed genes shorter than 300bp from the analysis, as they have a lower probability of insertions and are more likely to be falsely predicted to be essential (Table S11). Finally, we removed genes that had been deleted in the starting strains used for the mutagenesis, as no insertions in them would be detected.

**Fig. 3.**
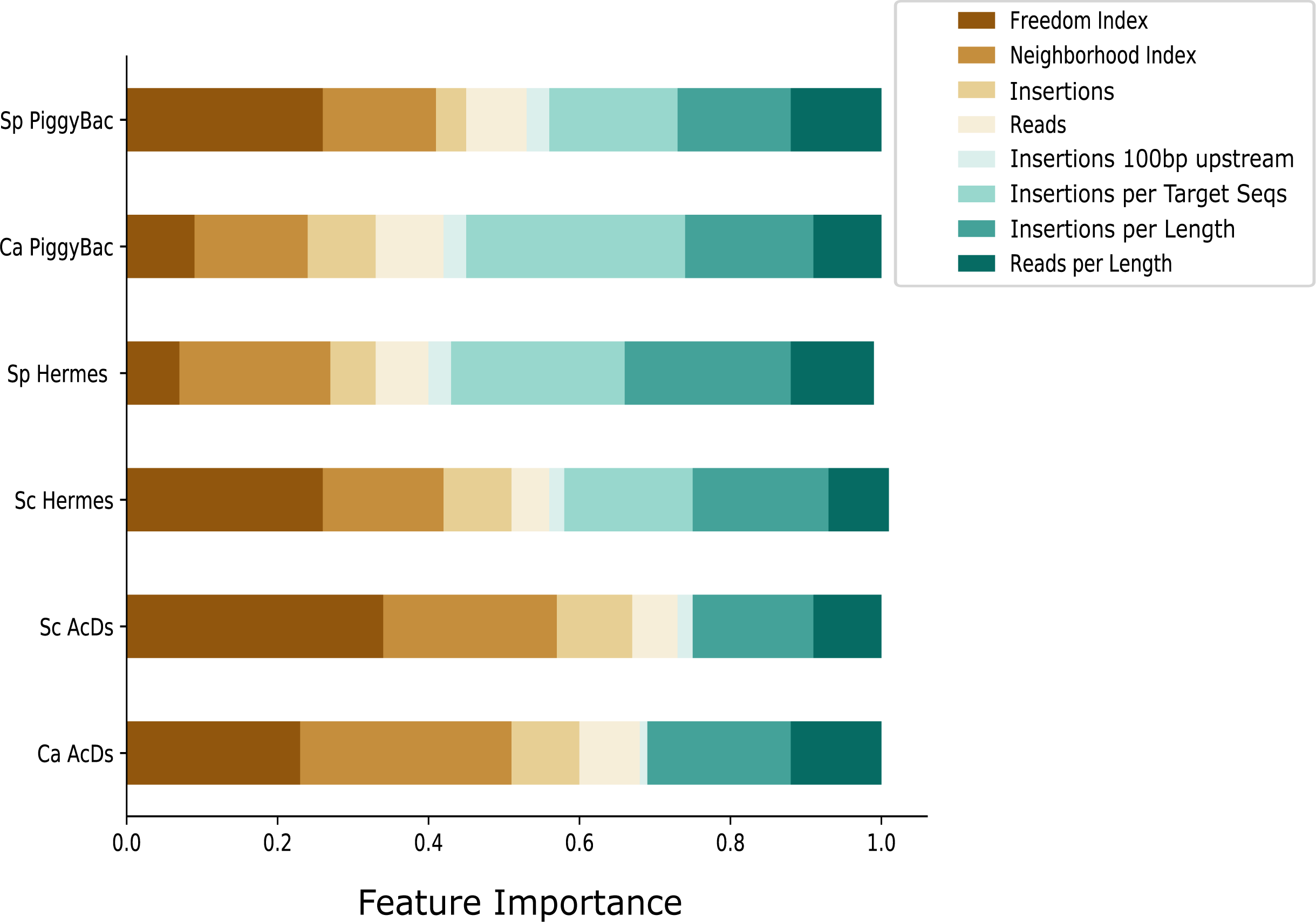
Comparison with known essentials genes. **a.** Comparison of the essentiality verdicts in *S. cerevisiae*, based on the known essential genes from the literature (SGD), *ScAcDs* (*Ac/Ds*) and *ScHermes* (*Hermes*) studies. **b.** Comparison of the essentiality verdicts in *S. pombe*, based on the known essential genes from the literature (PomBase), *SpPB* (*PiggyBac*) and *SpHermes* (*Hermes*) studies. **c.** Comparison of the essentiality verdicts in C. albicans, based on the known essential *S. pombe* and *S. cerevisiae* orthologs from the literature (Essential Orthologs), *CaAcDs* (*Ac/Ds*) and *CaPB* (*PiggyBac*) classifiers.

Classification thresholds differ slightly from the previously published analyses (Segal *et al*., 2018) based on threshold selection applied systematically to all 6 studies (described in detail in Methods).

Figures were generated using Python’s matplotlib and seaborn libraries (Bisong, 2019). The schematics were drawn using Inkscape (Bah, 2011). Mann-Whitney U p-values and Pearson’s correlation coefficients and p-values were calculated using Python’s Scipy (Millman and Aivazis, 2011).

## RESULTS AND DISCUSSION

### A comparative analysis of the transposon mutagenesis studies

We compared six *in vivo* transposon insertion mutagenesis experiments, produced using three different heterologous transposons (*AcDs, Hermes* and *PiggyBac*) in three different species (*S. cerevisiae, S. pombe and C. albicans*). Details of the datasets are provided in the methods section, and relevant parameters are highlighted in Table 2. The number of transposition events detected in the different studies varied considerably, from over 500,000 unique insertion sites (hits) and 84 million total reads for the *C. albicans AcDs* (*CaAcDs)*, to as few as 37,500 unique insertions and 6.1 million reads in the *S. pombe PiggyBac* (*SpPB*) data set (Table 2). The number of reads per insertion also varied considerably, from 41 to 170.

**Table 2.**
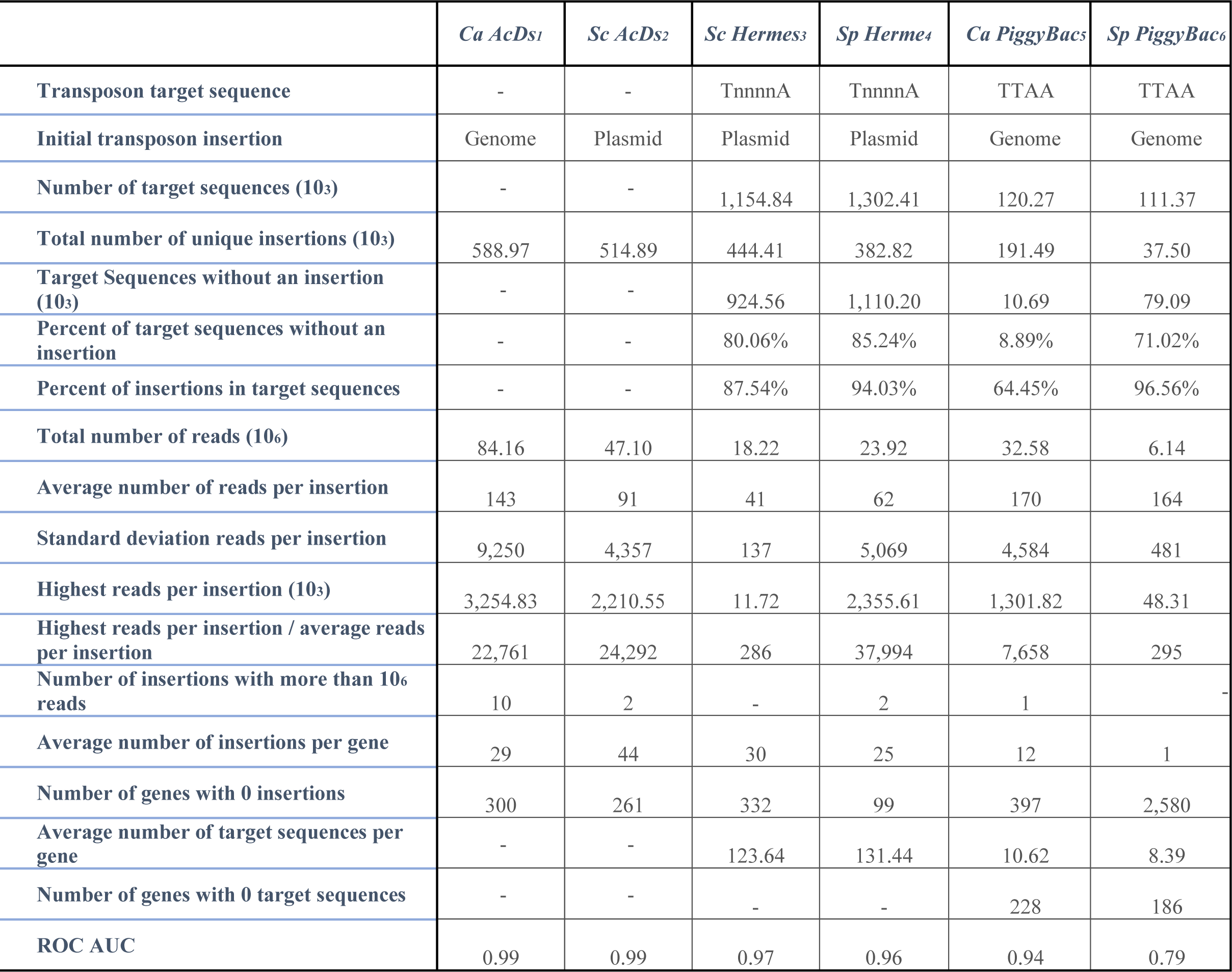
Summary of *in vivo* transposon mutagenesis libraries

### Overview of ML approach for gene essentiality prediction

We first mapped the insertion and read frequency of the transposons in each of the three reference genomes (Fig. 1c). We identified sites of transposon insertion based upon targeted sequencing of regions adjacent to the inserted transposon. Slightly different sequencing protocols were used in the different studies, but all six essentially amplified Tn-adjacent sequences and mapped them.

Theoretically, essential genes have no insertions and non-essential genes have many insertions; distinguishing essential and non-essential genes from the data, however, is not entirely straightforward (e.g., Fig. 1c, Gene X_n_). To address this ambiguity, we extended a previous approach for gene essentiality prediction(Segal *et al*., 2018). Briefly, we utilized a supervised learning approach (classification), designed to distinguish between different classes in the data (essential vs. non-essential) by learning the features of a previously labeled set of data points (training set; labeled as essential and non-essential genes, based on findings from deletion studies), and generalizing the predictions on the rest of the data (test set; genes unlabeled for essentiality). We chose Random Forest as the classifier, as its robustness is well established in genomic applications (Breiman, 2001; Chen and Ishwaran, 2012; Brieuc *et al*., 2018), and used the AUC (area under the receiver operating characteristic curve (Fig. 1d)) to analyze the specificity and sensitivity of the approach. Receiver operating characteristic is a graphical plot that illustrates the diagnostic ability of a binary classifier by plotting the true positive rate versus the false positive rate at various thresholds of classification (Hajian-Tilaki, 2013). The integral of this curve (area under the curve, AUC) is equal to the probability that the classifier will rank a randomly chosen positive instance (essential gene) higher than a randomly chosen negative instance (non-essential gene) or, in other words, the probability of a correct classification (Wray *et al*., 2010).

We first chose input features from transposon data that were likely to be informative in the essential/nonessential decision: the number of unique insertion sites (hits) per ORF, the degree to which those insertion sites were enriched in the population (reads), and the normalization factors that consider the insertion frequency as a function of chromosome position. We built the training sets (Table S1) using information from the two model species with gene essentiality data available from classical genetic approaches (e.g., comprehensive ORF deletion analysis) (reviewed in (Giaever and Nislow, 2014) and (Spirek *et al*., 2010)). For *C. albicans*, which did not have extensive prior knowledge of gene essentiality, we constructed a training set from a core set of genes whose orthologs were known to be essential in both model yeasts. This approach is likely to be useful for other species that lack sufficient prior knowledge of gene essentiality to construct a within-species training set.

We assessed the performance of each classifier by producing training sets using genes known or inferred to be essential and non-essential, as described in the Methods. Briefly, the classifiers were trained and their performance assessed using the AUC measure with a 5-fold cross-validation (training on 80% of the data and testing the performance on the remaining 20%, and averaging the 5 results). The AUCs were high across most examined studies (>0.94). The one exception was the *SpPB* study, which had far fewer unique insertion sites (Table 2) and an AUC of 0.785. The highest AUC levels were seen with the *AcDs* in both *C. albicans* and *S. cerevisiae*. Of note, these two studies also had the largest number of total insertions and reads. The considered ML features for each ORF in every study and the predicted verdicts of essentiality are provided in Tables S2-S7. Below, we describe the main insights gained from this comparison.

### Optimize the number of independent insertion sites (hits) for highest quality predictions of gene essentiality

The total number of unique insertion sites (hits) and the performance (AUC) values were highly correlated, and this correlation was statistically significant (Fig. 4a; Pearson’s r = 0.892; p-value = 0.0169). By contrast, the total number of sequencing reads showed a weaker correlation with the AUC that was not statistically significant (Fig. 4b; Pearson’s r = 0.636; p-value = 0.1741). If we disregard the worst performing *SpPB*, the correlation of the AUCs with the total number of insertions rises dramatically to Pearson’s r = 0.995; p-value = 0.0003, and the correlation with the total number of sequencing reads remains similarly weak: Pearson’s r = 0.652; p-value = 0.2327. A library with many independent insertions will thus improve performance, and simply increasing the number of sequencing reads will not likely be sufficient to obtain optimal results. Increasing the number of independent insertions requires collection of sufficient numbers of independent colonies soon after transposase induction and the resulting transposon excision and reinsertion. An advantage of *Hermes* is that most insertions occur during the stationary phase, so transposase-inducing conditions can be tolerated throughout the growth period. Experimental designs that optimize isolation of independent events are critical.

**Fig. 4.**
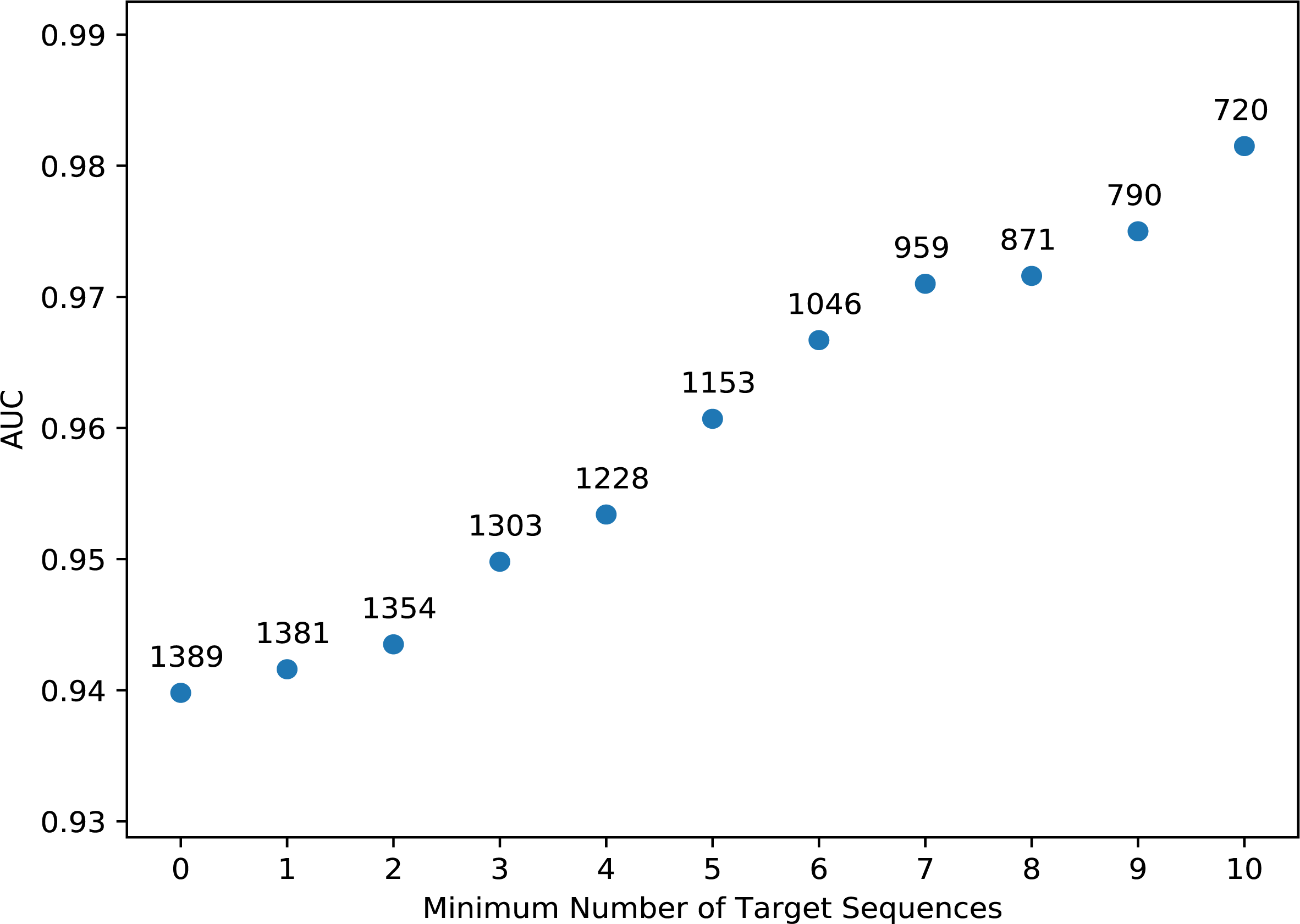
Contribution of unique insertions and total number of reads to the quality of ML predictions for gene essentiality/non-essentiality. The performance of the models (estimated in AUC) vs (**a**) the total number of unique insertion sites (hits) and (**b**) the total number of sequencing reads in each study (organism abbreviations as in Fig. 1; *Ca, C. albicans; Sp, S. pombe; Sc, S. cerevisiae*). Performance in each study was estimated using AUC (y-axis) and plotted against the total number of unique transposon insertion sites (10^3^), and the total number of sequencing reads (10^6^) obtained in each respective study.

### Avoid libraries with high levels of jackpot events

A jackpot event is the appearance of extraordinarily high numbers of reads in a very small number of insertion sites. When the number of reads greatly exceeds the theoretical number of cell divisions in the experiment, it is likely due to a transposition event happening prior to the induction of transposon excision in the experiment. Jackpot events are a major pitfall, with much sequencing capacity wasted on detection of a single insertion site. Jackpot events with >1M sequencing reads were present in 4 of the 6 data sets; *SpPB* and both *ScHermes* `studies had no major jackpot events (no insertion sites with ≥1000-fold more reads than the average read/insertion)` (Table 2). Both of these libraries also had far fewer total sequencing reads than the other studies (6M and 18M vs 24-84M for the other libraries).

Of note, within a data set, some individual experiments had far more jackpot events than others (Table 2), which would be expected if jackpot events occur stochastically. Importantly, these events were not clearly associated with one of the three species or with the transposon type. This suggests that jackpots arise from technical rather than biological issues.

It is important to avoid jackpot events because they reduce data quality considerably: the higher the number of reads at a few jackpot sites, the lower the number of informative insertions sites and reads. Avoiding the selection of cells in which a transposon was already mobilized is key to ensuring that the number of insertions and reads provides good genome coverage. Dividing the cultures into dozens of small cultures and then re-pooling these subcultures after transposase induction can effectively dilute out most jackpot events. Preparing several independent libraries and sampling a few sequences in each may also be worthwhile. If a tested library shows one sequence twice in one hundred colonies, for example, it is likely to be a >1M jackpot event.

### Consider which features are most important in the analysis of a given transposon

In decision-tree-based algorithms, such as Random Forest (Breiman, 2001), every node is a condition of a split of the data by a single feature. The splitting process continues until reaching a stop condition, such as: all the features being used, a very small obtained subset, or essentiality labels that are the same for the obtained subset. The goal is to reduce entropy (uncertainty) in the data. Entropy is zero when all labels in the obtained subset are the same; and is maximum when half of the labels in the obtained subset are the same (in a binary classification). Each split of the data by a given feature (node in the tree) reduces the entropy. The importance of a given feature in the Random Forest classifier is the calculated decrease in entropy contributed by that feature. Here we describe the features of the classifiers, and discuss their relative importance.

For each ORF, we catalogued predictive features including: the number of independent insertion sites, number of reads, and the length of each ORF. We also calculated engineered features including a neighborhood index, which normalizes for insertion bias due to genomic position (e.g., proximity to the initial excision site in the genome), and a freedom index, which reports the proportion of an ORF that is insertion-free (see Methods and Table 1). The neighborhood index is more important in studies where the transposon was originally placed in the chromosomal loci, but it also normalizes for any other insertional biases such as chromatin accessibility and 3D chromosome organization. The freedom index is especially useful for identifying genes with essential domains that are able to tolerate insertions outside of those domains. We used the number of transposon insertions per transposon target sequence in an ORF as an additional feature in the analysis (Fig. 5), where applicable (in *PB* and *Hermes* studies). We also calculated the number of insertions and the number of reads normalized by the length of each ORF, and compared the ‘feature importance’ for each library to ask whether specific features varied in importance for the inference of gene essentiality and whether feature importance was characteristic for a given transposon or yeast species.

**Fig. 5.**
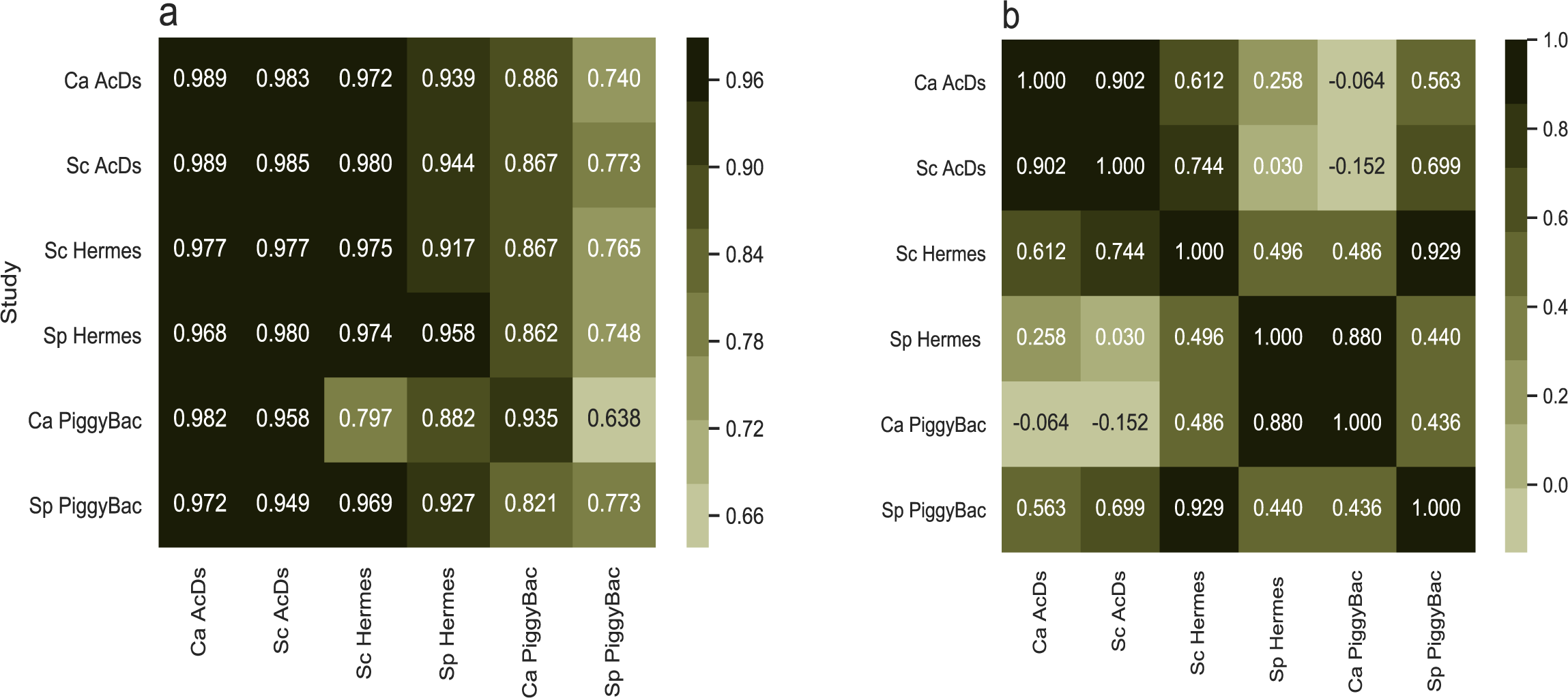
Feature importances in the different models of gene essentiality inference. Importance of each feature used in the Random Forest classifier of essentiality for each dataset. Features are described in Table 1; Neighborhood index generally normalizes for non-random insertion frequencies across the genome; Freedom index reports on the largest proportion of an ORF that has no insertions, which is a measure of domains that may be essential (Segal *et al*., 2018).

The number of insertions per ORF played an important role in determining essentiality, with essential genes having far fewer insertions than non-essential ORFs (∼7 times less, on average, across the 6 datasets), consistent with the strong correlation between number of insertions and the AUC (Fig. 4a). The number of reads per ORF played a lesser role in these classifications, also consistent with the correlation above (Fig. 4b). Gene length also affected the probability of transposon insertion in a gene, and thus was a crucial normalization parameter for the numbers of insertions and reads for every ORF.

The Neighborhood Index (NI) feature made important contributions in most of the classifications (except *SpPB*, which had far less data). Importantly, this feature did not differ considerably between the transposons induced from a plasmid and those inserted in the chromosomal loci (19.3 versus 19.6 respectively; Fig. 5). This result is consistent with the idea that chromatin accessibility, 3D chromosome organization, and other factors that may bias the insertion site frequency in a given organism can affect the frequency of insertion of different transposons in a similar manner. We attribute this phenomenon to the fact that while insertion of the transposon in a chromosomal loci creates a bias for nearby re-insertions, induction from a plasmid introduces other biases, such as preference for pericentromeric regions, as previously reported by (Michel *et al*., 2017).

The Freedom Index (FI) was a major contributor to both *ScAcDs* and *CaAcDs* predictions, while results with the *PB* and *Hermes* datasets were mixed (Fig. 5). This result is consistent with the idea that *AcDs* does not have a strongly specific target sequence and thus inserts throughout ORFs, and that *Hermes* have fewer target sequences within ORFs (on average, 123.65 per ORF in *S. cerevisiae* and 131.44 in *S. pombe*; Table 2), while *PB* has even fewer target sequences (on average, 10.62 per ORF in *C. albicans* and 8.39 in *S. pombe*; Table 2). Thus, the FI appears to be more important in *AcDs* experiments because insertions occur more randomly throughout an ORF.

The importance of the number of insertions in the proximal regulatory sequences (100 bp upstream to the start codon) to the essential/non-essential predictions was only minor, but highly variable. For example, the impact of both *ScHermes* studies was nearly twice that of *ScAcDs* for this feature. As described in more detail below, we posit that this difference is due to cryptic enhancer/promoter activity in the *miniDs* transposon in *S. cerevisiae* not seen with either one of the *Hermes* transposon (Fig. 6). Surprisingly, this cryptic promoter doesn’t seem to negatively affect the inference of gene essentiality, as seen in the higher performance of the models in *Ac/Ds* studies (AUCs, Table 2; Fig.2) relative to studies with different transposons, *CaAcDs*, however, is slightly better in this regard than *ScAcDs*.

**Fig. 6.**
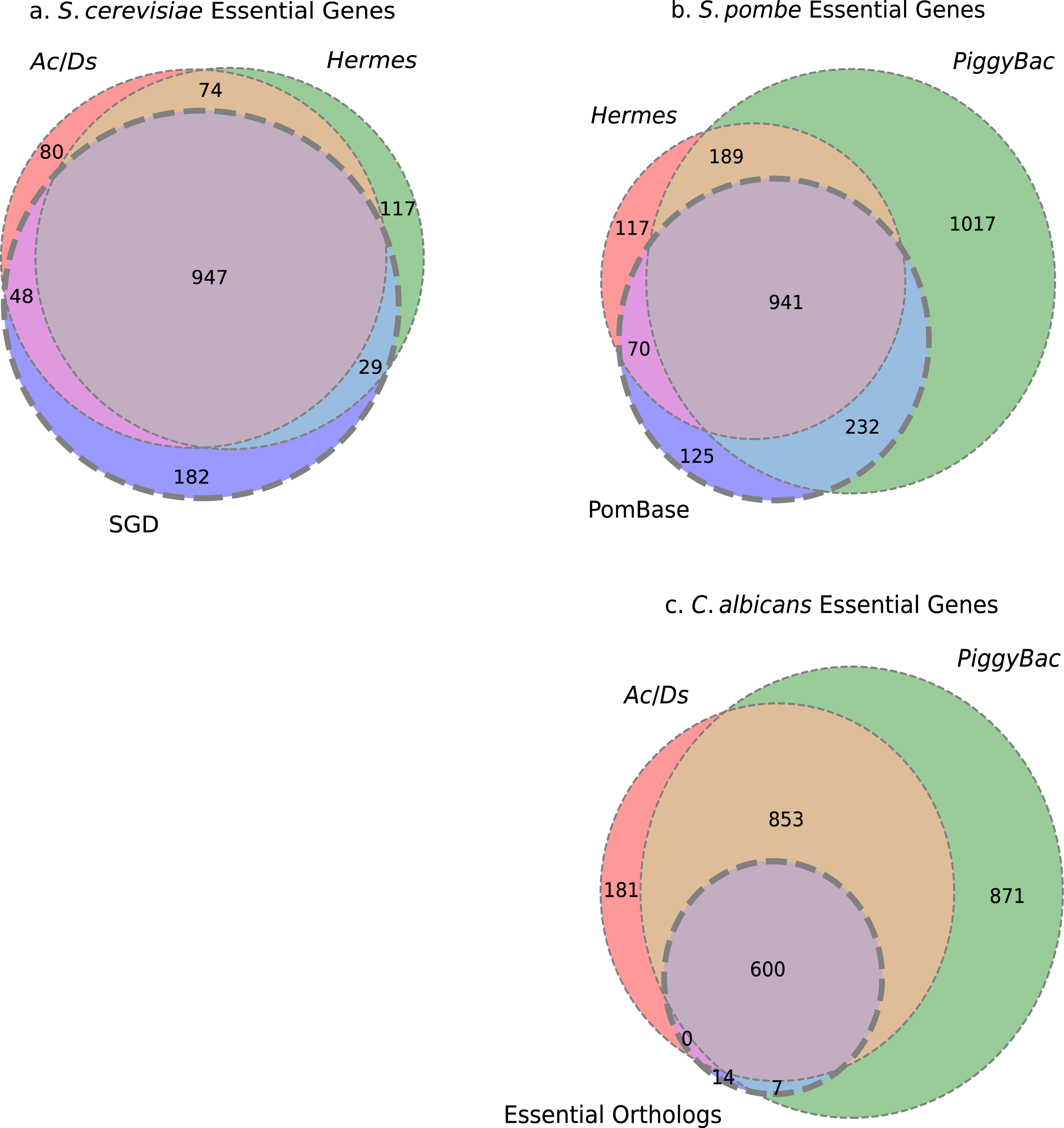
*Hermes* and *AcDs* transposon insertion distribution in *S. cerevisiae* ORFs. Transposon insertions surrounding the start codons of *S. cerevisiae* ORFs were plotted for non-essential (**a**) and essential genes (**b**). *Hermes* in a haploid strain (blue), *Hermes* in a diploid strain (black) and *AcDs* (*miniDS*, red). X-axis-coordinates (bp) relative to the start codon; Y-axis-number of insertions per site per gene (10^8^). In non-essential genes (**a**), the insertion distribution patterns appear to be indistinguishable in the three studies. In contrast, essential genes (**b**) are more tolerant to insertions in the diploid strain than in haploids. Furthermore, haploid strains tolerated *AcDs* insertions up to 35bp upstream to the start codon, while they did not tolerate *Hermes* insertions within 200bp upstream to the start codon.

### Consider the effect of transposon-specific target sequence specificity

Most transposons have preferred sites of insertion: *Hermes* prefers TnnnnA (Supplementary Figures S13-14) and *PiggyBac* inserts primarily at TTAA sequences (Supplementary Figures S15-16); *AcDs*, in contrast, does not have a strong insertion site preference (Supplementary Figures S11-12). Theoretically, the length of the insertion site sequence necessarily scales inversely with the number of potential unique insertion sites. It was not clear, however, at what insertion sequence length the resolution of studies of gene essentiality becomes limiting.

The feature importance of the number of insertions per transposon target sequence in an ORF, which should be a measure of library saturation, showed a varying degree of importance in the *PB* and *Hermes* studies (Fig. 5). It was much more important in *PB* than *Hermes* studies, especially if we disregard the lower quality classification of *SpBP* (0.29 vs 0.17 and 0.23, respectively). It should be noted that the target sequences are preferred sites of insertion, yet are not exclusive or absolute (logo analysis; Supplementary Figures S11-16). For example, *PiggyBac* in *C. albicans* had 1.6-fold more unique insertion sites than the theoretical number of target sequences in the *C. albicans* genome. By contrast, for both *ScHermes* and *SpHermes*, the number of target sequences available far outnumbered the number of unique insertions. The proportion of target sequences without insertions ranged from 8.9% for *CaPB* to 85% for *SpHermes* (Table 2) and the proportion of insertions not in target sequences ranged from 3.3% in *SpPB* to 36.6% in *CaPB*, and from 12.5% in *ScHermes* to 6.0% in *SpHermes*. We thus surmise that there are sufficient numbers of target sequences for *Ac/Ds* and *Hermes* as to not be a limiting factor for these transposons. This is not the case in *PiggyBac*, which doesn’t seem to have enough target sequences throughout the yeasts’ genomes.

Another critical issue is the number of genes that lack any preferred target sequences within the ORF; there are 228 and 185 ORFs without a single TTAA sequence in *C. albicans* and *S. pombe*, respectively. These ORFS have a lower probability of acquiring insertions and, if the genes are non-essential, are much more likely to give false positive information (be predicted essential for lack of insertions). Indeed, 155 ORFs without TTAA sequences were predicted essential in the *CaPB* data and yet predicted non-essential in the *CaAcDs* study. Similarly, 118 of the 185 ORFs lacking TTAA sequences were predicted essential from the *SpPB* study, but were non-essential in the *SpHermes* study. We assume that many of these genes are false positives, especially given that 127 of the 185 ORFs lacking TTAA, including 95 of the 118 aforementioned ORFs, were non-essential in classical genetics studies of *S. pombe*.

Next, we asked if the number of target sequences within an ORF affected the *CaPB* classification performance for that ORF. To address this, we compared the performance (AUC) to sets of ORFs filtered to exclude ORFs with different numbers of target sequences (from 0 to 10) from the training set used to train the classifier (Fig. 7). The AUC increased from ∼0.94 for the entire training set to >0.98 for the training set containing only genes with 10 or more target sites (∼50% of the genes in the training set). This suggests that studies using the *PiggyBac* transposon may struggle to correctly infer gene essentiality for ORFs with low numbers of target sites.

**Fig. 7.**
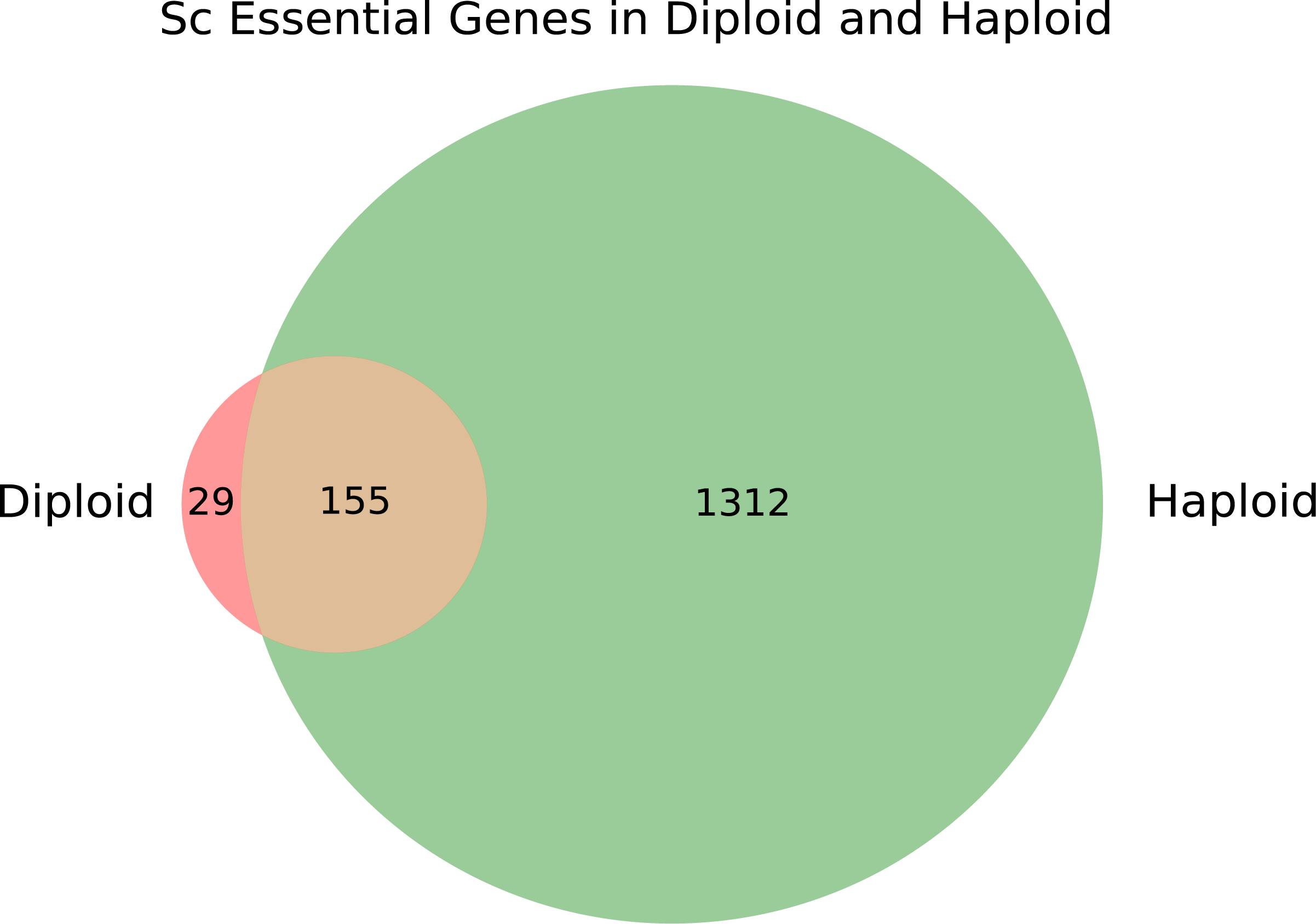
Analysis of the ability of the *CaPB* classifier to infer gene essentiality in genes with increasing number of target sequences. When only ORFs with a specific number of target sites are considered (>= x-axis), AUC rises accordingly (blue dots), but the number of ORFs that can be analyzed necessarily decreases (numbers above blue dots). This demonstrates the importance of the prevalence of the transposon target sequences in ORFs, for the quality of gene essentiality inference, using in-vivo transposon mutagenesis studies. X-axis: Minimum number of target sequences per ORF needed for inclusion in the classification. Y-axis: *CaPB* classifier AUC.

### Consider whether the transposon can activate as well as disrupt gene expression

Predictions of gene essentiality were based upon the assumption that transposon insertion into an ORF disrupted gene expression and produced loss-of-function allele. However, this is not necessarily the case for all genes. If an insertion allele removes a regulatory domain from a protein, for example, the protein may become hyperactive, resulting in a gain-of-function allele. Additionally, some transposons may introduce enhancer and promoter activities that could increase gene expression in some species. The *miniDs* transposon used in the *ScAcDs* data is likely to contain such activities (Michel *et al*., 2017). Consistent with this idea, the *ScAcDs* dataset contains an average of 4.71 insertions within the first 100 bp of essential genes (divided by the total number of insertions and multiplied by 10^6^), whereas the other datasets— including *CaAcDs*, which has a transposon modified from the *miniDs—* contain significantly fewer (1.98 insertions in the first 100 bp of essential genes). *SpHermes* and *ScHermes* contain 2.68 and 3.53 insertions (2.56 from (Edskes et al. 2018)), respectively, within the first 100 bp of essential genes. Additionally, while many essential genes of *S. cerevisiae* appeared to tolerate *miniDs* insertions, this is not true for *Hermes* insertions at sites in the 5’ UTR that are very close to the start codon (Fig. 6). The *miniDs* transposon in *S. cerevisiae* may thus facilitate inappropriate activation of gene expression when inserted upstream or within certain genes, as previously reported by (Michel *et al*., 2017).

### Cross-study analysis

Knowing the full set of essential and non-essential genes in eukaryotic microbes, including pathogens of humans, animals, and plants, will improve the understanding of common and species-specific properties of these understudied organisms. Furthermore, once a transposon library has been collected, it can be screened under many other growth conditions to reveal genotype/phenotype relationships. *In vivo* transposon analysis of gene essentiality is a practical and feasible approach, because the cost in time and resources for obtaining libraries is far lower than that for producing engineered deletion mutants, especially given that the amount of baseline information (other than the genome sequence) about the organisms may be minimal. The only technical hurdle is to introduce the heterologous transposon of interest, either on a plasmid (where feasible) or into a useful locus within the genome of the relevant organism.

An additional challenge is that ML approaches require a high-quality training dataset of gold-standard essential and non-essential genes. For many non-model organisms, such training data is too sparse to build a robust training set. For *C. albicans*, we circumvented the low numbers of genes already known to be essential by relying upon genes that had been determined to be essential from comprehensive classical genetic deletion studies in both model yeasts (*S. cerevisiae* and *S. pombe)* and those with orthologs in *C. albicans*. Training on *S. cerevisiae* or *S. pombe* orthologs with consistently essential orthologs yielded good performance predictions for *C. albicans* (AUC: 0.940 to 0.993, Table 2). *CaAcDs* performance was lower when training only on the 66 genes known to be essential plus the set of presumed non-essential genes (those that had been successfully deleted in *C. albicans* studies, AUC of ∼0.92) (Segal *et al*., 2018).

Next, we considered the quality of the learning performance for each dataset if we trained on orthologs from one species and predicted essentiality of genes in a different organism (Fig. 8). The orthologous training set was designed to have a high overlap of essential genes between the *S. cerevisiae* and *S. pombe* training sets; out of the 721 *S. cerevisiae* essential genes, 621 were also essential in *S. pombe* and all of the *S. pombe* essential genes were essential in *S. cerevisiae*. For each cross-study examination, we trained a model on 80% of the training set of the organism in the training study and tested the performance on 20% of the labeled genes of the organism in the target study (Fig. 8a). The transfer learning performance of the classifications was of a comparable quality to the single study classifiers for most *AcDs* and *Hermes* cases (Fig. 8a and Fig. 4). Furthermore, it displayed a somewhat symmetrical property: in most cases, there were minor differences in performance between train/test and test/train pairs (reducing the quality by ∼0.5% to 5.7%) when the tests were between or among *AcDs* and *Hermes* experiments. Conversely, when testing for predictions from *PB* data that were trained on either *AcDs* or *Hermes*, the AUCs dropped more dramatically (up to ∼21.9%). *PB* data was thus less transferable than *Hermes* and *AcDs* data.

**Fig. 8.**
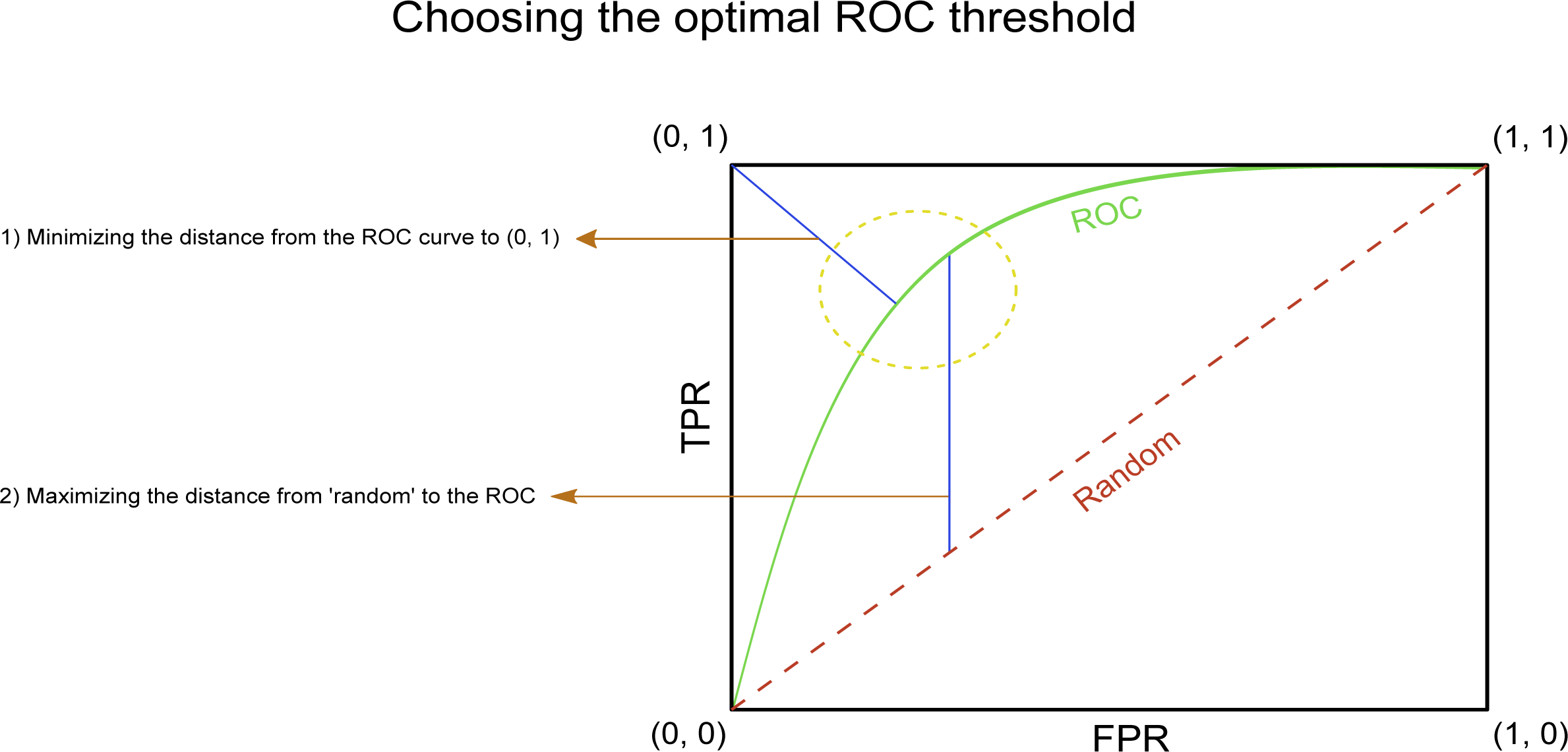
Analysis of model performance, trained on data from a different organism and/or transposon in all possible combinations. **a.** For each ROC AUC value in the table, training was performed on 80% of the original training set used in the training species/transposon described in the rows. This training data was then used to predict the essentiality of the remaining 20% of the training set in the species/transposon described in the columns. The train/test split ratio was similar to the 5-fold cross-validation performed in the single study analyses. **b.** For each study, the vector of the relative feature importance was correlated with the feature importance in all the other studies. Pearson r correlation coefficient values are presented.

The low *PB* transferability between *SpPB* and *CaPB* is likely due to the sparser target sequence distribution relative to either *Hermes* or *AcDs*, which causes *PB* studies to produce false positives, as noted above, and thus might contribute to reduced performance in cross-study analyses. The lower performance of the classifiers in the original *PB* single studies (Table 2) also may have contributed to the reduced ability to predict essentiality in pools of *PB* mutants using cross-species models.

Reduced differences in cross-study performance could also be due to similarities between the importance of different features in the classifiers for the different datasets. To test this possibility, we correlated the vector of the relative feature importance for each study with the feature importance in all the other studies (Fig. 8b). The analysis distinguished 3 groups within the 6 studies, based on the correlation coefficient values for feature importance between members of the group: *CaAcDs* and *ScAcDs* (Pearson’s r = 0.902); *Sp Hermes* and *CaPB* (Pearson’s r = 0.898); and *SpPB* and *ScHermes* (Pearson’s r = 0.929). Notably, with the exception of the *AcDs* studies, the quality of the transfer learning predictions appears to be independent of both the transposon type and the organism studied. We presume that this is due to the lack of a specific target sequence for the *AcDs* transposon system.

### As necessary, construct training sets using genes with orthologs in models where essentiality is known and then validate the training set manually

We suggest that an initial training set of orthologous genes known to be essential and non-essential in related model organisms can be used to facilitate analysis of a transposon insertion study in a non-model organism with sparse essentiality information (Table S8). An important caveat is that differences between gene function in different species can alter gene essentiality of a small number of these orthologs; it is thus important to visually inspect this orthologous training set before applying it. The goal is to remove any genes with insertion patterns that are highly contradictory to the ‘essentiality label’ that the orthologs provided. For example, for *C. albicans*, three independent inspectors reviewed the entire orthologous training set in an unprejudiced fashion, visually examining the insertion patterns in the *CaAcDs* data and manually labeling each gene as essential, non-essential, or ambiguous. When all three inspectors classified a gene as non-essential (e.g., many insertions throughout an ORF within a genome region that had many insertions outside of that ORF) and the orthologs were labeled ‘essential’ in the two model yeasts, we removed that gene from the training set.

Once a training set has been established, and the features for the ORFs have been calculated, the Random Forest classifier can be run in a cross-validation scheme and the AUC can be calculated using the essentiality labels. This provides an efficient approach to obtaining information about all the genes in a species that has been sequenced but not subjected to much molecular manipulation. Clearly, the same approach can be used to compare the essentiality of the same sets of genes grown in different conditions as well, potentially providing large amounts of phenotypic data across an entire set of ORFs. If applied to a species that has not been the subject of many genetic studies, such data would represent a treasure-trove of information about the genes themselves and the phenotypes associated with loss-of-function of those genes.

### A comparative analysis of gene essentiality predictions

One important issue is whether different transposon insertion studies in the same organism have similar or different predictions from one another and from the known essentiality status of deletion mutants, which are by definition ‘loss-of-function’ null alleles. After removing genes that were repeated or <300 nt in length (see Methods), 74 genes were predicted to be essential by both *S. cerevisiae* transposon studies and were listed as non-essential by SGD (Fig. 3a). A recent study of the *S. cerevisiae* deletion (YKO) collection used whole genome sequencing to identify genome changes that may improve the growth of different deletion mutants (Puddu *et al*., 2019). Indeed, 29 of the 74 genes that were essential in both Tn studies and non-essential in SGD, carried at least one genomic aberration (e.g. aneuploidy, copy number variation of rDNA, telomeric length, copy number variation of mDNA and etc.) (Table S9). Such genomic aberrations may compensate for reduced competitive fitness of the specific deletion mutants, and it is presumed that such aberrations were selected in the course of conventional strain construction steps (Puddu *et al*., 2019). Because random *in vivo* transposon mutagenesis involves far less selective pressure than the construction and selection of specific deletion mutants, Tn insertion mutant strains are less likely to have accumulated such putative compensatory mutations. Accordingly, the isolates carrying insertions in these genes may have had lower competitive fitness that was interpreted as essentiality. The fact that insertions in these genes were rare in both *S. cerevisiae* transposon studies supports this hypothesis. Of the remaining 45 genes, 34 were not in TKO collection sequenced by (Puddu *et al*., 2019), presumably because of their poor growth and low fitness. Consistent with this, most of these genes are annotated in SGD as having reduced competitive fitness, making them more likely to be at low frequency in the population of strains carrying transposon insertions. Furthermore, strains tested for essentiality in the YKO collection were tested on medium buffered to neutral pH, while the Tn selection conditions were on media that are considerably more acidic (pH4.5-5). Neutral pH likely facilitates the growth of some mutants, such as several <u>v</u>acuolar membrane <u>a</u>tpase (VMA) genes that were essential in the Tn studies but not in the YKO collection. It is also important to note that gene essentiality, although treated as a binary or discrete function (essential or not), is actually a common quantitative trait not only in *S. cerevisiae* (*Liu et al*., *2015*), but also in *S. pombe* (Li *et al*., 2019). We posit that many of the genes determined to be essential in both transposon studies but not in the YKO collection are likely to be genes with traits that exhibit quantitative essentiality and/or are suppressible by genome aberrations that accumulated in the YKO collection (Puddu *et al*., 2019) but not in the Tn insertion mutants.

For *S. cerevisiae*, the classifiers displayed a high degree of agreement on the final verdicts of gene essentiality (Fig. 3a), while more discrepancies were evident for the *C. albicans* and *S. pombe* studies. Both *PiggyBac* studies predicted a much higher number of essential genes than the *AcDs* or *Hermes* studies (Fig. 3b and 7c), as expected from the paucity of target sequences and the lower number of total unique insertions which, as discussed above, make false positive predictions more likely. For example, the *CaPB* study had an average 5.84 target sites per kb in genes likely to be false positives vs. 10.62 target sites per kb in all the genes (Mann Whitney U: p-value < 2.38*e^- 78^). Importantly, when compared to the set of essential genes for each species that were determined by deletion analyses, the transposon studies also did quite well, with only 20 to 35% of the genes in disagreement. In some cases, such discrepancies were found to be due to issues with the deletion collection isolates. For example, ∼8% of the original *S. cerevisiae* deletion collection carried aneuploidies or gene amplifications (Hughes *et al*., 2000), and ∼10% of *S. pombe* deletion strains retained a wild-type copy of the ORF that had been targeted for deletion. Extra copies of the ‘deleted’ gene reduce the apparent number of essential genes. Furthermore, differences in culturing conditions and experimental protocols introduce another source for misdiagnosis, as mutations that reduce fitness or are conditionally essential, might be called as essential. Due to the aforementioned discrepancies, we suggest carefully evaluating the essentiality calls given by this method, and relying on data from more than one independent study or source. In this case, we can only recommend genes that were considered essential from two independent transposon mutagenesis studies. Previously, we formulated a more stringent approach, which considered transposon mutagenesis results and several independent deletion studies (Segal *et al*., 2018).

### Gene essentiality in haploid versus diploid strains of *S. cerevisiae*

*S. cerevisiae* is readily grown in both the diploid and haploid states, allowing identification of the haplo-insufficient subset of genes among the set of essential genes. Based on gene knockout studies, we classified only 2 genes (*NDC1, MLC1*) as haplo-insufficient (Stevens and Davis, 1998; Chial *et al*., 1999), and all other essential genes as haplo-proficient (i.e. heterozygous knockouts in diploids were viable). To determine whether additional haplo-insufficient genes exist in *S. cerevisiae*, we collected *ScHermes* insertions in diploid strain BY4743 and compared them to haploid strains BY4741 and BY4742. We inferred gene essentiality in *ScHermes* transposon mutagenesis libraries using the model that had been trained on the *SpHermes* haploid training set data, applying the same threshold for classification (Fig. 9, Fig. 2). The classifier identified 155 genes as ‘essential in both haploid and diploid’, a number far higher than expected. Upon closer analysis, 98 contained regions of poor mapping due to duplications elsewhere in the genome, 50 were categorized as dubious in the Saccharomyces Genome Database (*Saccharomyces Genome Database*, no date), one (*LEU2*) had been deleted in the strains studied, and the two known haploinsufficient genes (*NDC1, MLC1*) had been identified, providing support for this approach. Upon visual inspection of data of the remaining four genes, one essential gene (*BCY1*) appeared haploinsufficient, whereas another (*RPC10*) contained numerous insertions in its 5’ UTR in diploids but not haploids, suggesting that it might not be essential (Fig. 10). Although there are

∼50% more insertions in the diploid study, the discrepancy in the number of insertions becomes clearly evident only in genes that were classified as essential in the haploid studies (Fig. 6). The other two ORFs predicted to be haploinsufficient are very small (165-225 bp) and within regions of sparse insertion density. We thus have lower confidence in the data for these two genes. Transposon mutagenesis of a diploid strain successfully revealed the two known haploinsufficient genes and inferred a new one (*BCY1*), which is known to be essential in the conditions employed in the screen but not essential in other culture conditions (Matsumoto *et al*., 1982). The haploinsufficiency of *BCY1* should be validated experimentally.

**Fig. 9.**
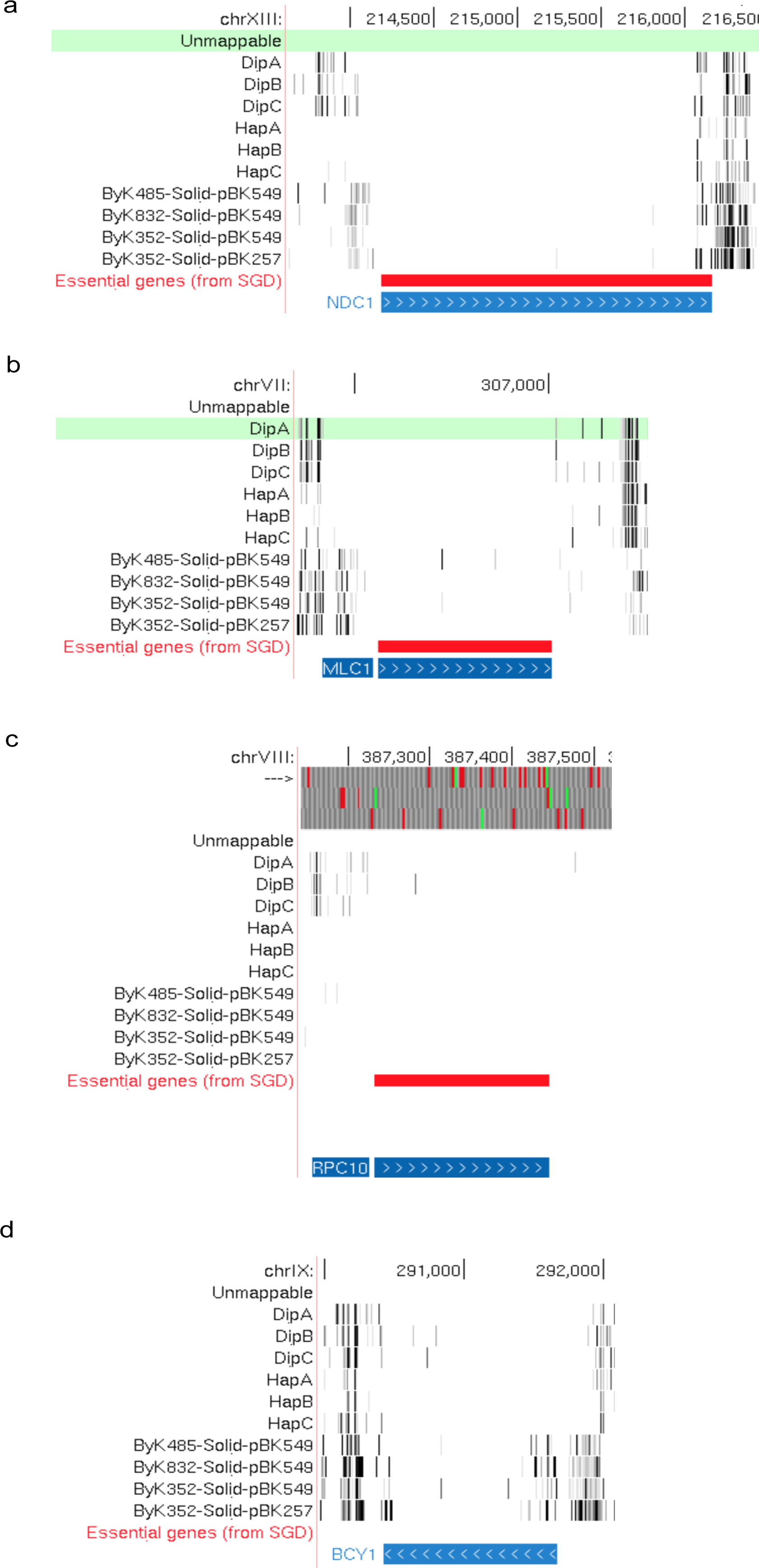
Gene essentiality in haploid and diploid *S. cerevisiae*. Comparison of essential genes in haploid and diploid *S. cerevisiae* analyzed with *ScHermes*. Random Forest classifier was trained on the haploid *ScHermes* study and predicted gene essentiality in a diploid strain, using the same threshold for the final verdict. Mitochondrial genes were not considered.

**Fig. 10.**
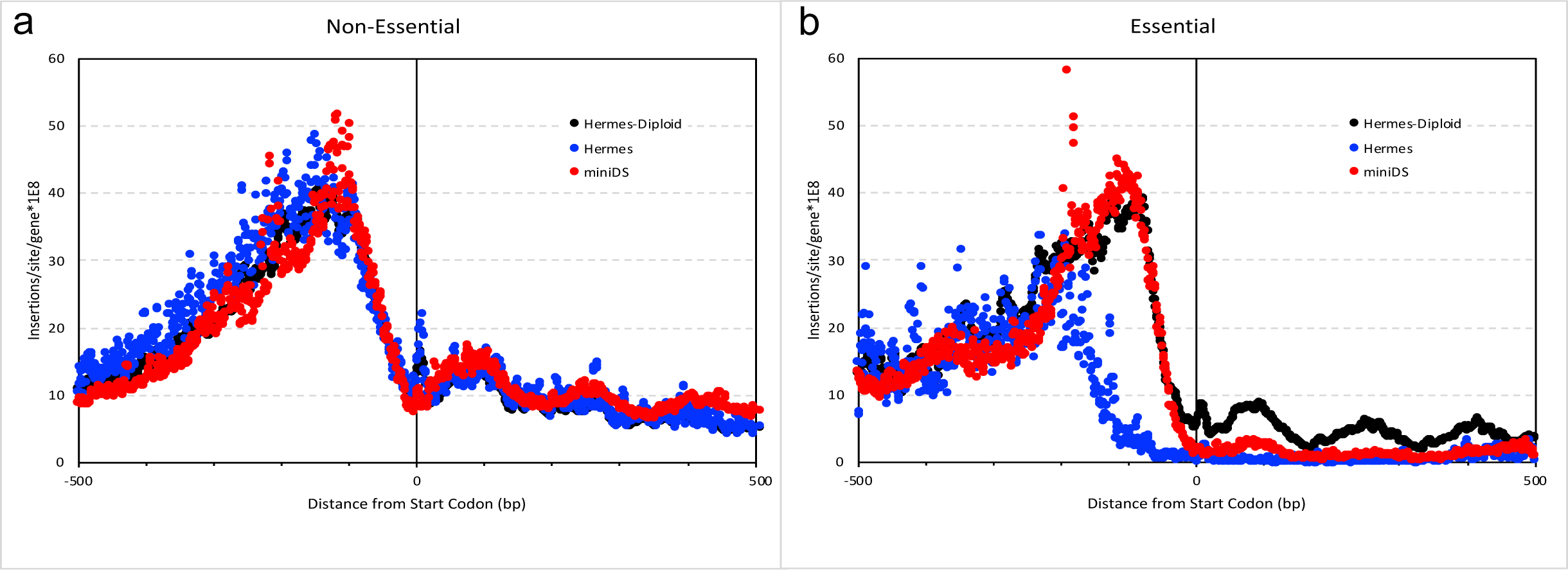
Suspected haploinsufficient genes in *S. cerevisiae*. Four genes suspected to be haploinsufficient in *S. cerevisiae: NDC1, MLC1, RPC10 and BCY1*, as they appear in the UCSD genome browser (‘UCSC Genome Browser on S. cerevisiae Apr. 2011 (SacCer_Apr2011/sacCer3) Assembly’, no date). *NDC1* (**a**) and *MLC1* (**b**) were known to be haploinsufficient. *BCY1* (**d**) appears to be a previously unknown haploinsufficient gene. *RPC10* (**c**) might not be essential as it sustained insertions in the 5’ UTR in diploids but not haploids.

The classifier also identified 29 genes as ‘haploinsufficient’ in diploids and not essential in haploids. Of these, 20 could be dismissed based on their annotation as dubious or the presence of duplicated (unmappable) segments. All of the remaining 9 genes were small (87-528 bp) and found in regions of sparse insertion density. These genes are annotated in SGD as non-essential and are probably false positives.

These observations raise an important issue about data quality control. It is important to filter dubious and uninformative ORFs from the data set (as was done in the analysis of *CaAcDs* (Segal *et al*., 2018)). This includes genes with repeated domains or duplicate copies in the genome that prevent unambiguous mapping of short Illumina reads. Furthermore, predicting the essential/non-essential status for short ORFs, and especially those located in regions with sparse intergenic insertions, is more likely to be problematic.

## CONCLUSIONS

**In summary**, we suggest a number of metrics for the inference of gene essentiality using *in vivo* transposon mutagenesis studies in yeasts, including those with little available genetic data. Maximizing the total number of unique transposon insertions is the most critical factor in achieving optimal performance of the classification. This can be attained by collecting many independent insertion clones, striving to reduce possible jackpot events in the study, maximizing the depth of coverage, and utilizing a transposon with a fairly permissive target sequence or no preferred target sequence. Furthermore, while transposons with relatively stringent target sequences have some advantages for screens that identify individual mutants, they are less robust for determining gene essentiality. The low number of potential target sequences, and especially the lack of any target sequences, in certain ORFs will increase the likelihood of falsely classifying non-essential genes as essential (Fig. 7). Additionally, we think that transposon mutagenesis is an ideal approach for gaining large amounts of useful genotype/phenotype data about understudied organisms; the cross-species learning methodology allows inference of gene essentiality based on conserved orthologs, especially when coupled with visual screening of the data. Finally, *in vivo* transposon mutagenesis is an incredibly useful tool for high throughput genomic studies, not only of gene essentiality *per se*, but also of genes required under specific selective conditions. We hope that the recommendations provided here will facilitate future work to understand genes in a wide range of yeast species.

## DECLARATIONS

## FUNDING ACKNOWLEDGEMENTS

European Research Council Advanced Award 340087 (RAPLODAPT) to J.B., the Israel Science Foundation grants no. 715/18, 757/12 to R.S. and no 997/18 to J.B. and by NIH R21-AI130722 and T32-GM007231 to KWC.

A.L. is supported by a fellowship from the Edmond J. Safra Center for Bioinformatics.

## Author Contribution

A.L. designed the study with J.B. and R.S., E. K. D., D.W.K., A.B.G. and K.W.C. acquired the *S. cerevisiae Hermes* data and commented on its analysis; A.L. analyzed these new data, as well as the published transposon datasets with input from J.B. and R.S.. A.L. and J.B. wrote the manuscript, A.L., R.S. and J.B. edited the manuscript.

## Competing Interests

The authors declare no competing interests.

## Data Availability

The *S. cerevisiae Hermes* data (mapped reads and counts) are available at http://genome-euro.ucsc.edu/s/CunninghamLab/Hermes%20Vs%20AcDs. The code is available at https://github.com/berman-lab.

## Supplementary Information

Supplementary Figures S1-S17 are attached separately with their associated legends, in a single PDF file. Supplementary Tables S1-S11 are attached in separate CSV files.

^1^ (Segal *et al*., 2018)

^2^ (Michel *et al*., 2017)

^3^ Data from (Edskes *et al*., 2018) was included in Table S4b, and compared to the other *S. cerevisiae* data in Fig. S17

^4^ (Guo *et al*., 2013)

^5^ (Gao *et al*., 2018)

^6^ (Li *et al*., 2011)

## Notes

### Competing Interest Statement

The authors have declared no competing interest.

### Summary of Updates

Analysis was redone to exclude false positives that were very small ORFS and duplicated genes. New figures 9, 10 and 11 and revisions to Figure 7.

