## Supplementary Material for "Comparing the utility of *in vivo* transposon mutagenesis approaches in yeast species to infer gene essentiality"

The following tables are attached in separate CSV files:

Table S1 Training Sets.
Sets of essential and non-essential genes used as training sets for the inference of gene essentiality.

Table S2 Machine Learning features and predictions of gene essentiality in *C. albicans AcDs.*
A table with all the features considered for the classification and the predictions of essentiality of each ORF in the *Ca AcDs* study

Table S3 Machine Learning features and predictions of gene essentiality in *S. cerevisiae AcDs.*
A table with all the features considered for the classification and the predictions of essentiality of each ORF in the *Sc AcDs* study

Table S4a Machine Learning features and predictions of gene essentiality in *S. cerevisiae Hermes.*
A table with all the features considered for the classification and the predictions of essentiality of each ORF in the *Sc Hermes* study

Table S4b Machine Learning features and predictions of gene essentiality in *S. cerevisiae Hermes* from Edskes *et al*., 2018.
A table with all the features considered for the classification and the predictions of essentiality of each ORF in this *Sc Hermes* study

Table S5 Machine Learning features and predictions of gene essentiality in *S. pombe Hermes.*
A table with all the features considered for the classification and the predictions of essentiality of each ORF in the *Sp Hermes* study

Table S6 Machine Learning features and predictions of gene essentiality in *C. albicans PiggyBac.*
A table with all the features considered for the classification and the predictions of essentiality of each ORF in the *Ca PB* study

Table S7 Machine Learning features and predictions of gene essentiality in *S. pombe PiggyBac.*
A table with all the features considered for the classification and the predictions of essentiality of each ORF in the *Sp PB* study

Table S8 Orthologs with known essentiality.
A set of all *S. pombe* and *S. cerevisiae* orthologous to *C. albicans* with their essentiality

Table S9 74 Disagreeing *S. cerevisiae* genes.
List of 74 *S. cerevisiae* genes that were deemed essential by both transposon studies and non-essential by SGD

Table S10 Lists of genes with duplications.
Lists of genes with paralogs in *S. cerevisiae, C. albicans* and *S. pombe*

Table S11 Lists of genes shorter than 300bp.
Lists of genes shorter than 300bp in *S. cerevisiae, C. albicans* and *S. pombe*

Fig. S1 Histograms of reads distribution in the *C. albicans AcDs* study.
A histogram of read distribution throughout the genome is provided in the *Ca AcDs* study. The y-axis is the number of reads on a logarithmic scale and the x-axis is the coordinate in the genome (chromosomes are sequentially appended one to another).

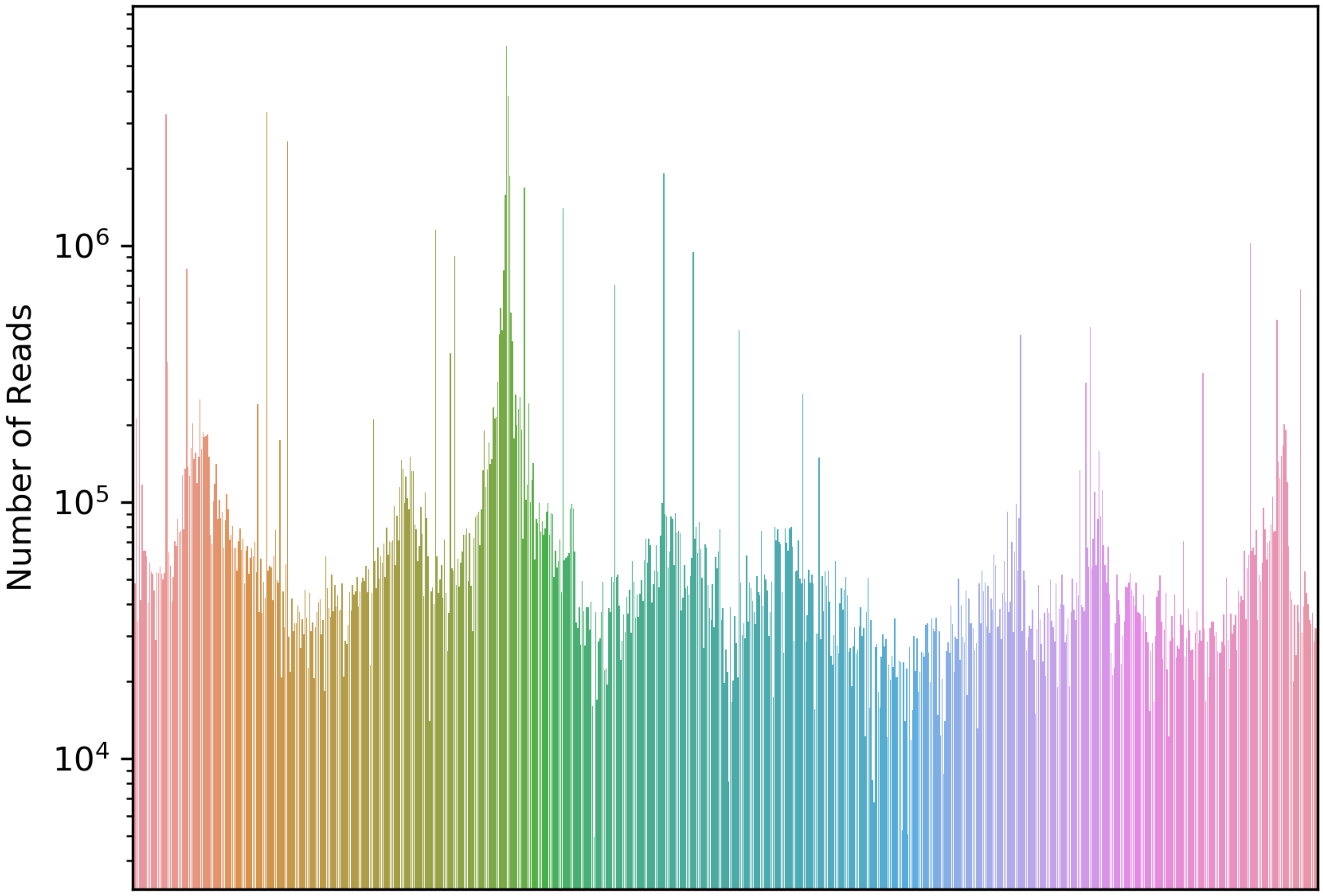

Fig. S2 Histograms of reads distribution in the *S. cerevisiae AcDs* study.
A histogram of read distribution throughout the genome is provided in the *Sc AcDs* study. The y-axis is the number of reads on a logarithmic scale and the x-axis is the coordinate in the genome (chromosomes are sequentially appended one to another).

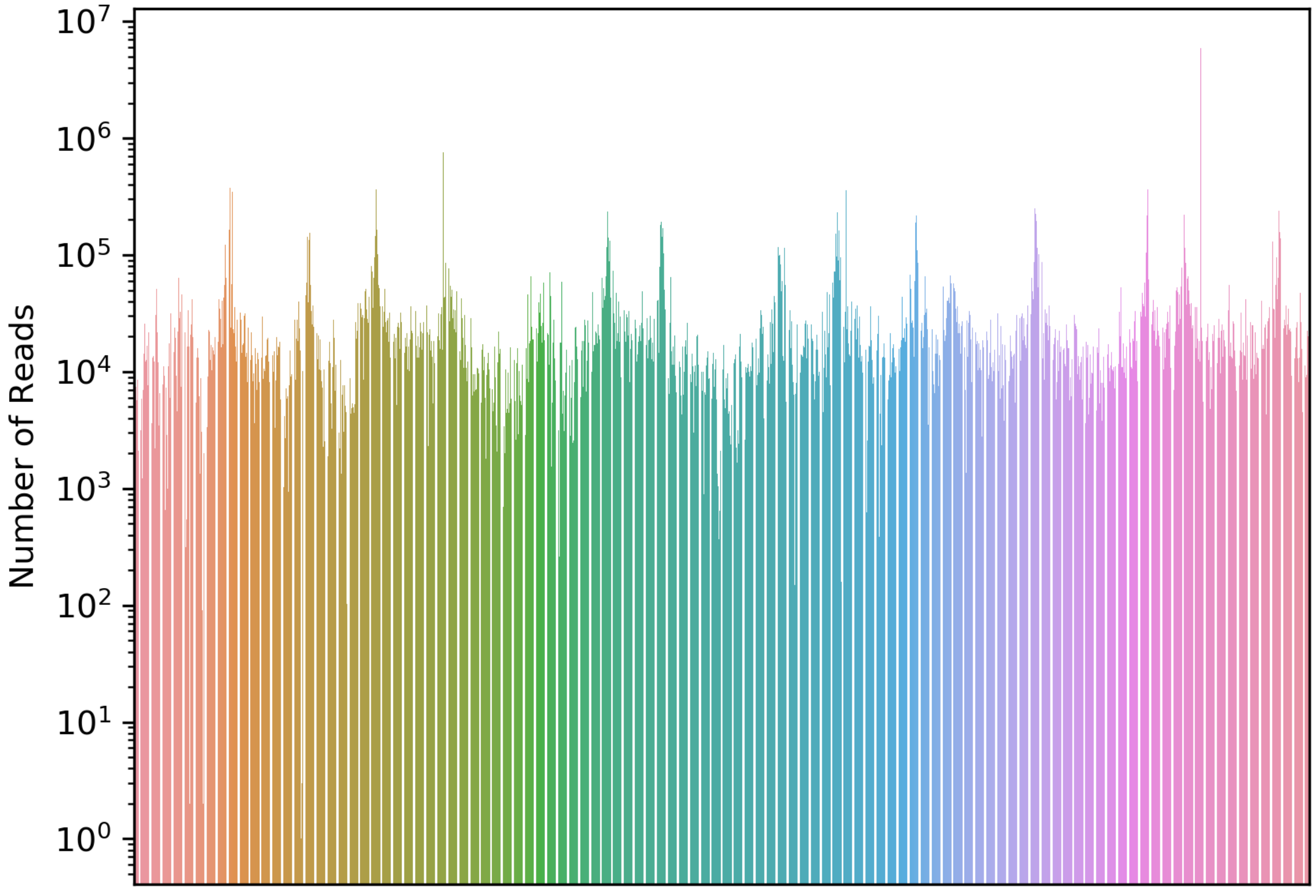

Fig. S3 Histograms of reads distribution in the *S. cerevisiae Hermes* study.
A histogram of read distribution throughout the genome is provided in the *Sc Hermes* study. The y-axis is the number of reads on a logarithmic scale and the x-axis is the coordinate in the genome (chromosomes are sequentially appended one to another).

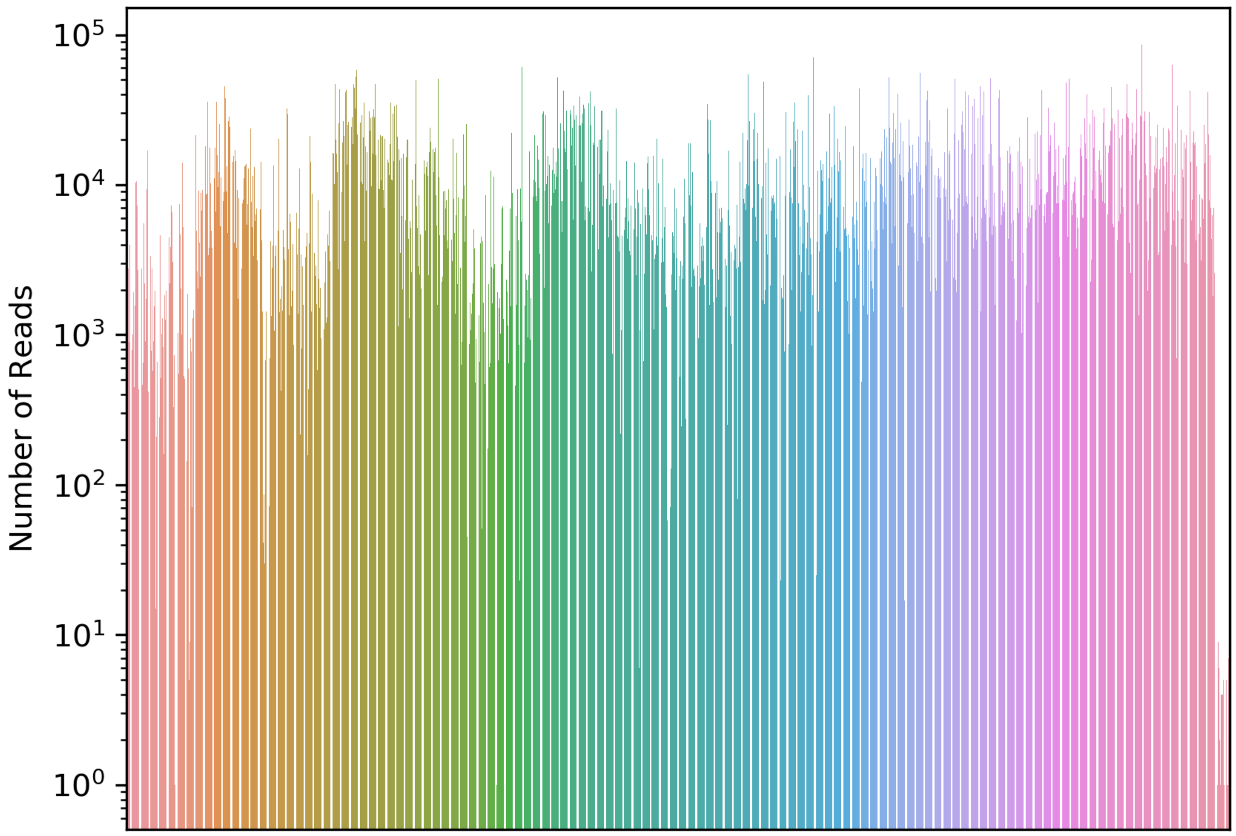

Fig. S4 Histograms of reads distribution in the *S. pombe Hermes* study.
A histogram of read distribution throughout the genome is provided in the *Sp Hermes* study. The y-axis is the number of reads on a logarithmic scale and the x-axis is the coordinate in the genome (chromosomes are sequentially appended one to another).

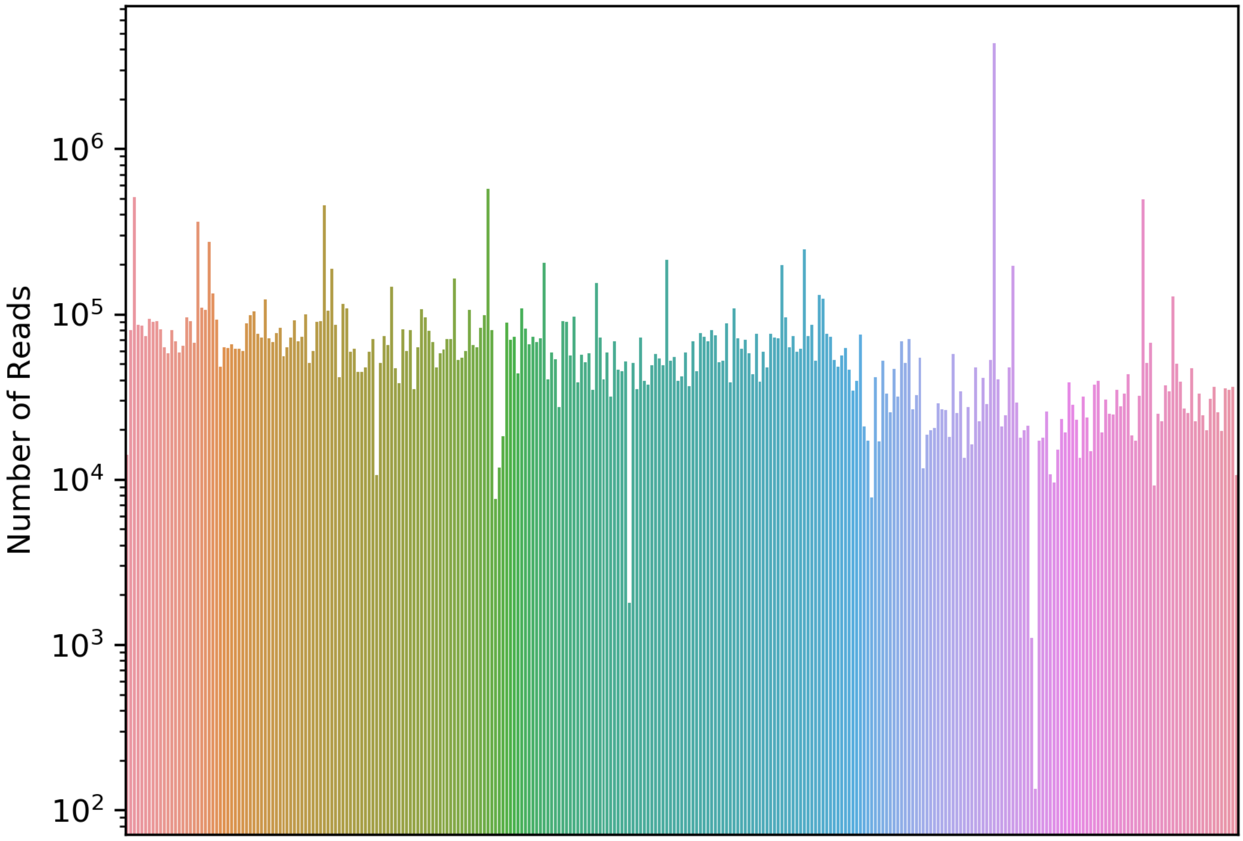

Fig. S5 Histograms of reads distribution in the *S. pombe PiggyBac* study.
A histogram of read distribution throughout the genome is provided in the *Sp PB* study. The y-axis is the number of reads on a logarithmic scale and the x-axis is the coordinate in the genome (chromosomes are sequentially appended one to another).

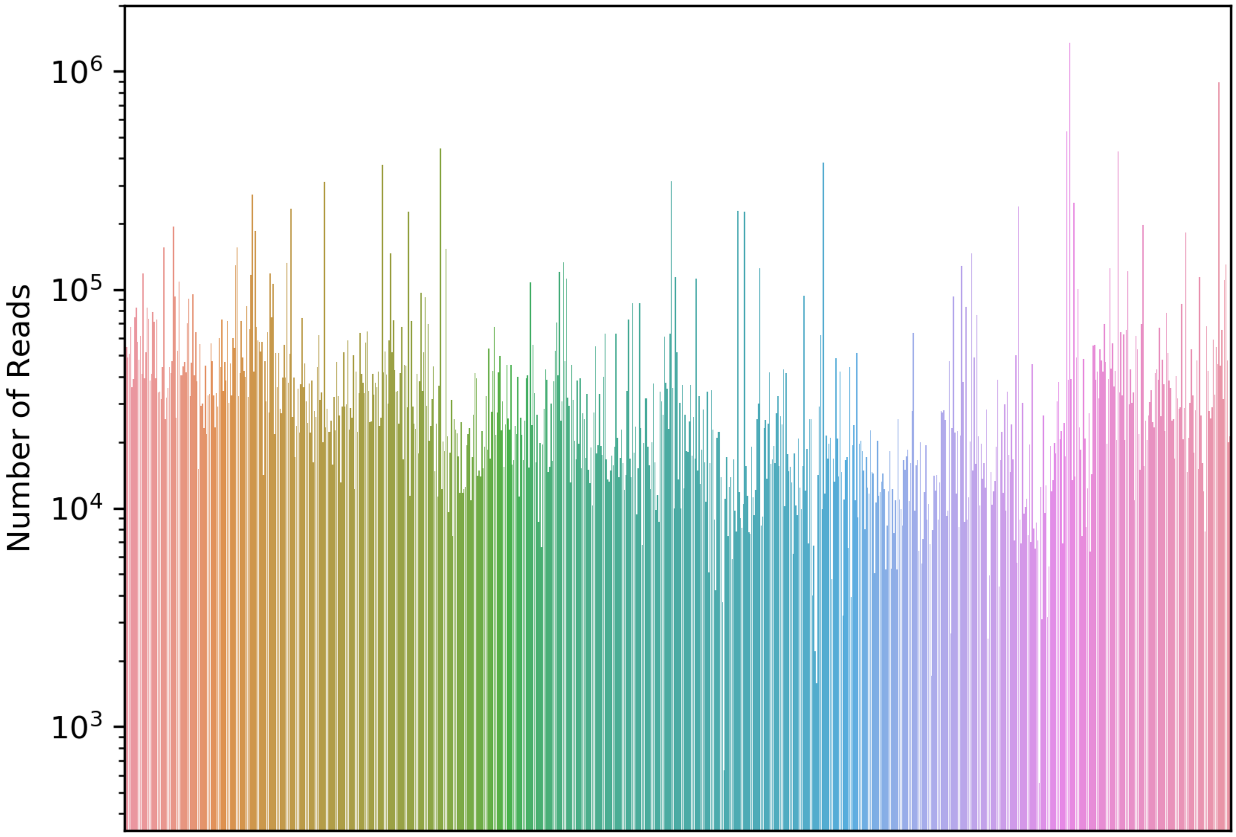

Fig. S6 Histograms of reads distribution in the *C. albicans PiggyBac* study.
A histogram of read distribution throughout the genome is provided in the *Ca PB* study. The y-axis is the number of reads on a logarithmic scale and the x-axis is the coordinate in the genome (chromosomes are sequentially appended one to another).

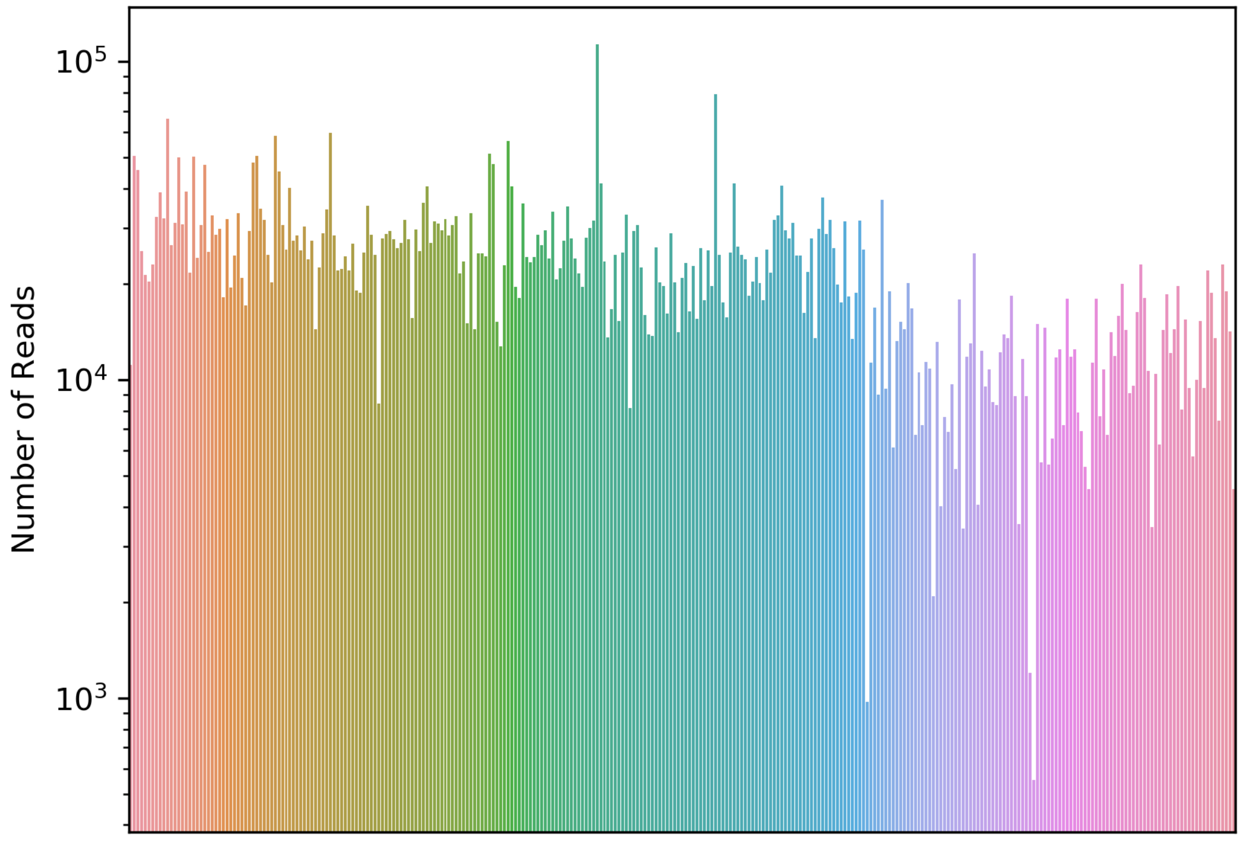

Fig.s S7 Frequency of transposon target sequence at the location of the transposon insertions in the *S. cerevisiae Hermes* study.

A histogram of the frequency of transposon target sequence at the location of the transposon insertions is provided for the *Sc Hermes* study. The y-axis is the percentage of finding a transposon target sequence and the x-axis is the location relative to all the transposon insertions.

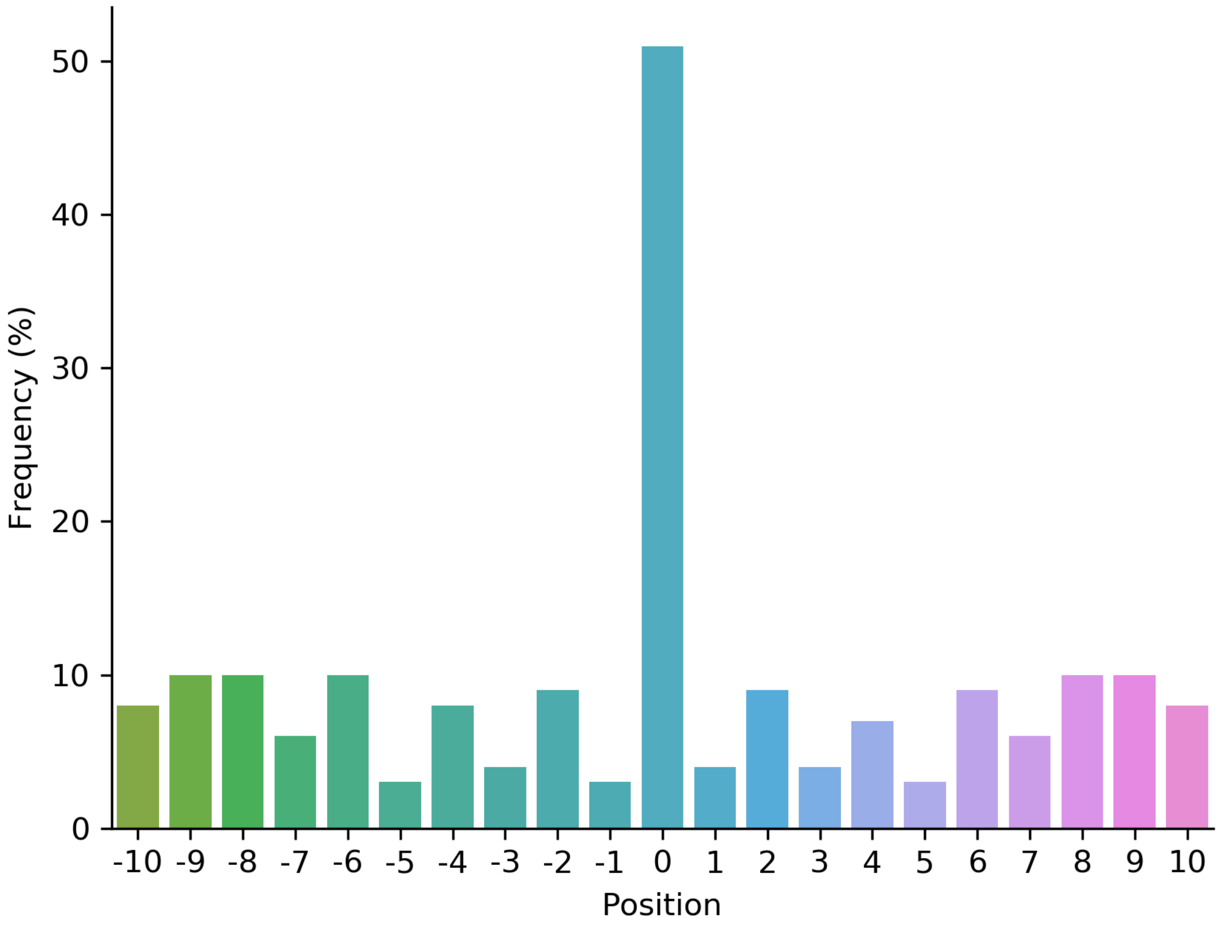

Fig. S8 Frequency of transposon target sequence at the location of the transposon insertions in the *S. pombe Hermes* study.
A histogram of the frequency of transposon target sequence at the location of the transposon insertions is provided for the *Sc Hermes* study. The y-axis is the percentage of finding a transposon target sequence and the x-axis is the location relative to all the transposon insertions.

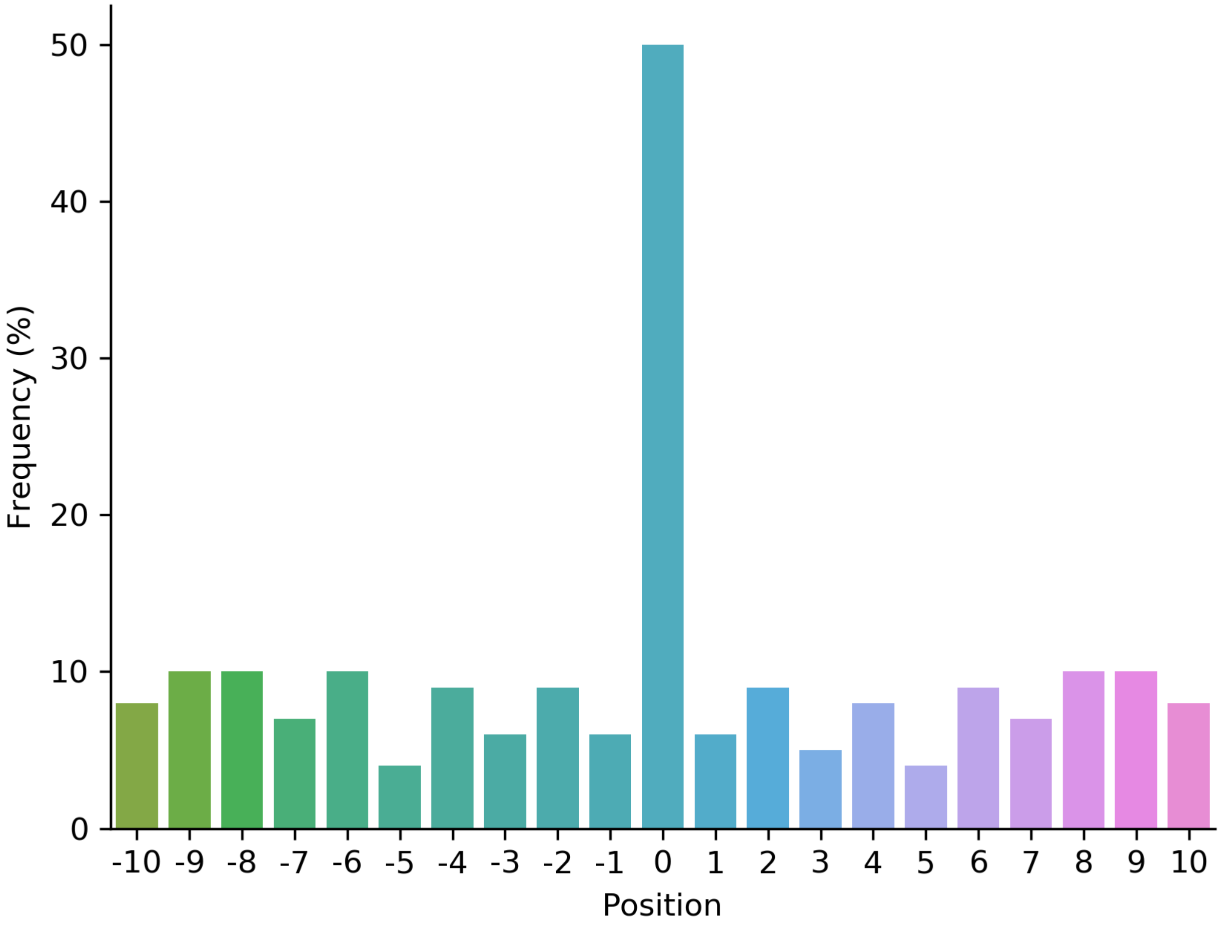

Fig. S9 Frequency of transposon target sequence at the location of the transposon insertions in the *C. albicans PiggyBac* study.
A histogram of the frequency of transposon target sequence at the location of the transposon insertions is provided for the *Ca PB*. The y-axis is the percentage of finding a transposon target sequence and the x-axis is the location relative to all the transposon insertions.

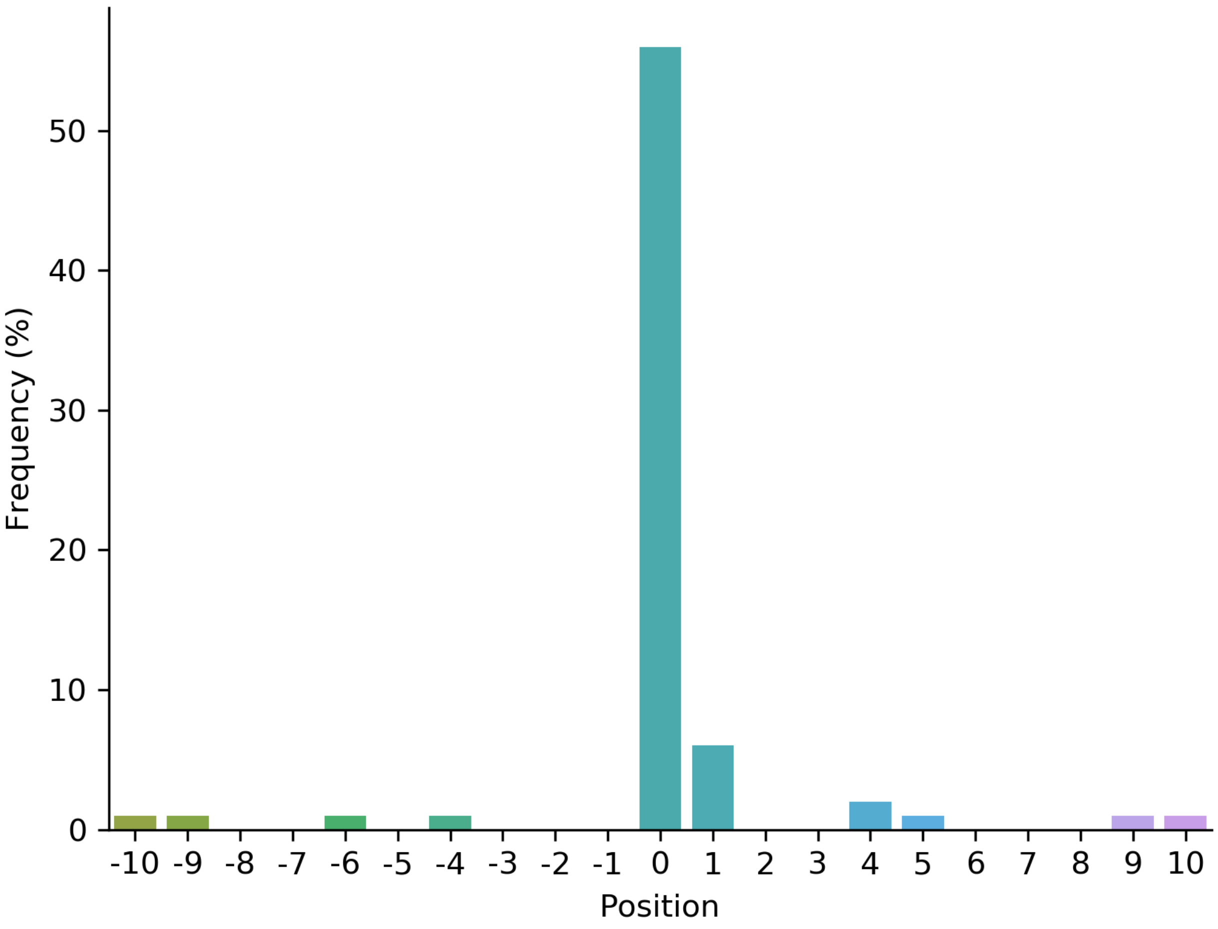

Fig. S10 Frequency of transposon target sequence at the location of the transposon insertions in the *S. pombe PiggyBac* study.

A histogram of the frequency of transposon target sequence at the location of the transposon insertions is provided for the *Sp PiggyBac* study. The y-axis is the percentage of finding a transposon target sequence and the x-axis is the location relative to all the transposon insertions.

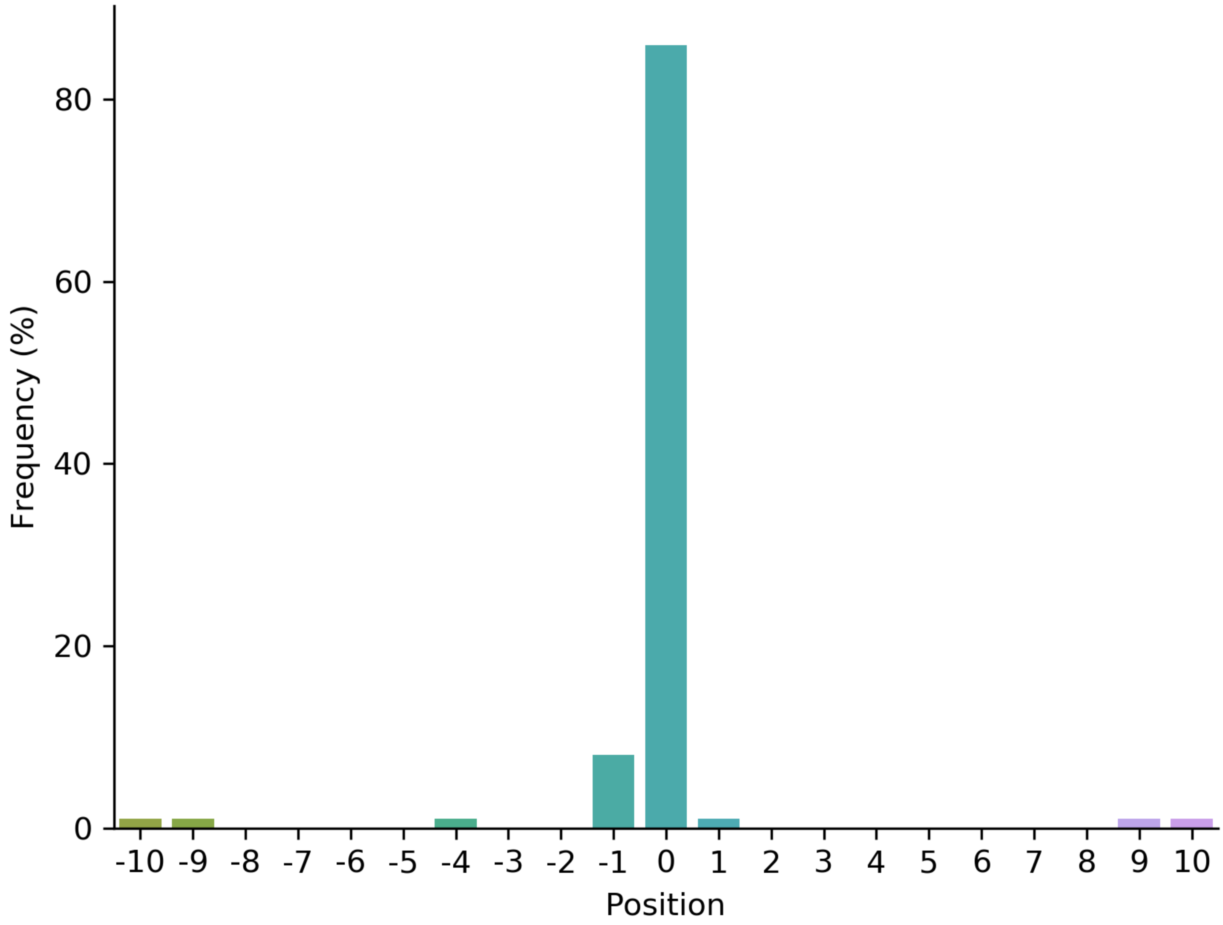

Fig. S11 Logo motif analysis of *ScAcDs* study.
A graphic representation of the frequency of nucleotides surrounding the *AcDs* transposon insertions in the *S. cerevisiae* genome

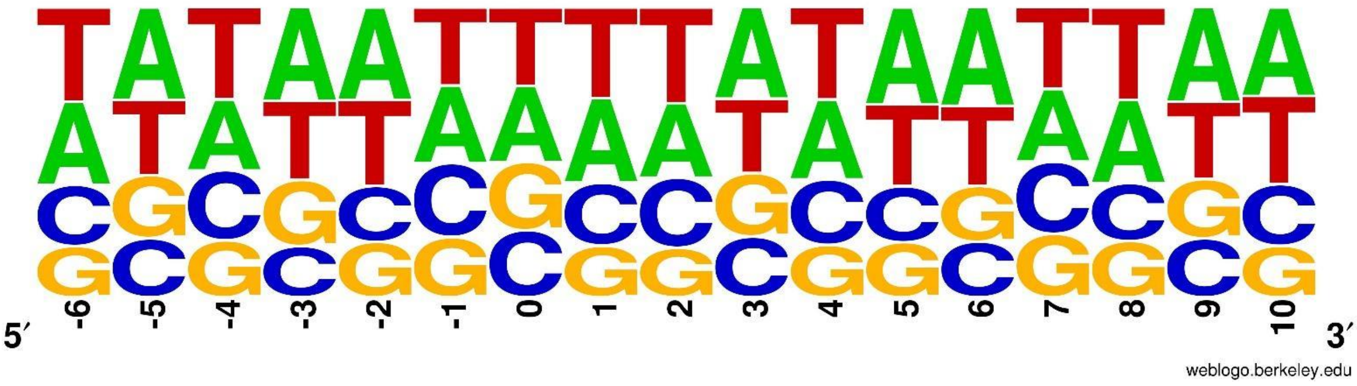


Fig. S12 Logo motif analysis of *CaAcDs* study.
A graphic representation of the frequency of nucleotides surrounding the *AcDs* transposon insertions in the *C. albicans* genome

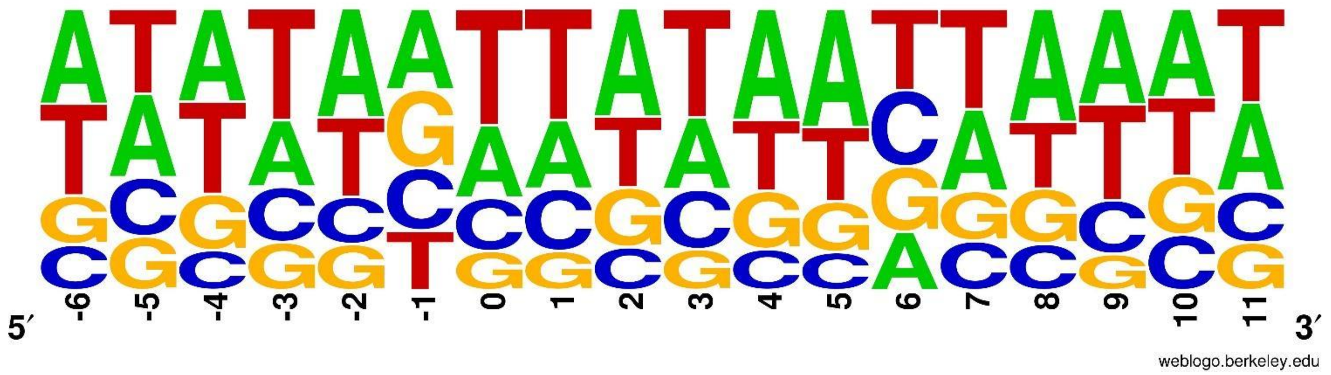


Fig. S13 Logo motif analysis of *ScHermes* study.
A graphic representation of the frequency of nucleotides surrounding the *Hermes* transposon insertions in the *S. cerevisiae* genome

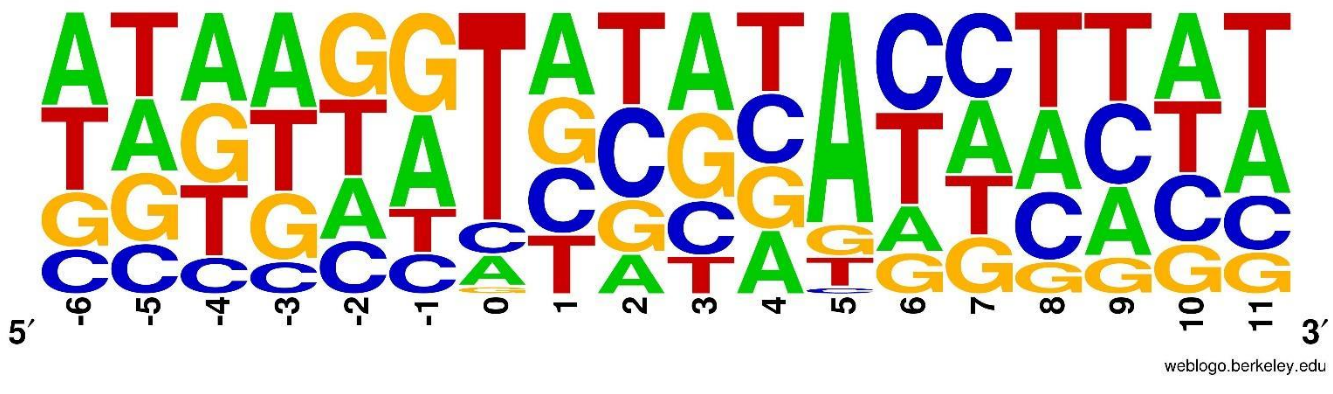

Fig. S14 Logo motif analysis of *SpHermes* study.
A graphic representation of the frequency of nucleotides surrounding the *Hermes* transposon insertions in the *S. pombe* genome

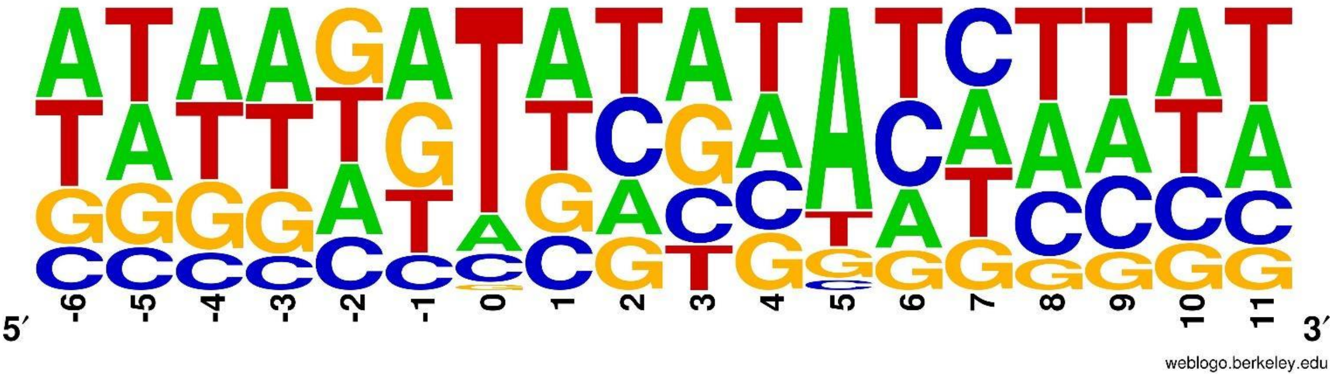


Fig. S15 Logo motif analysis of *SpPB* study.
A graphic representation of the frequency of nucleotides surrounding the *PB* transposon insertions in the *S. pombe* genome

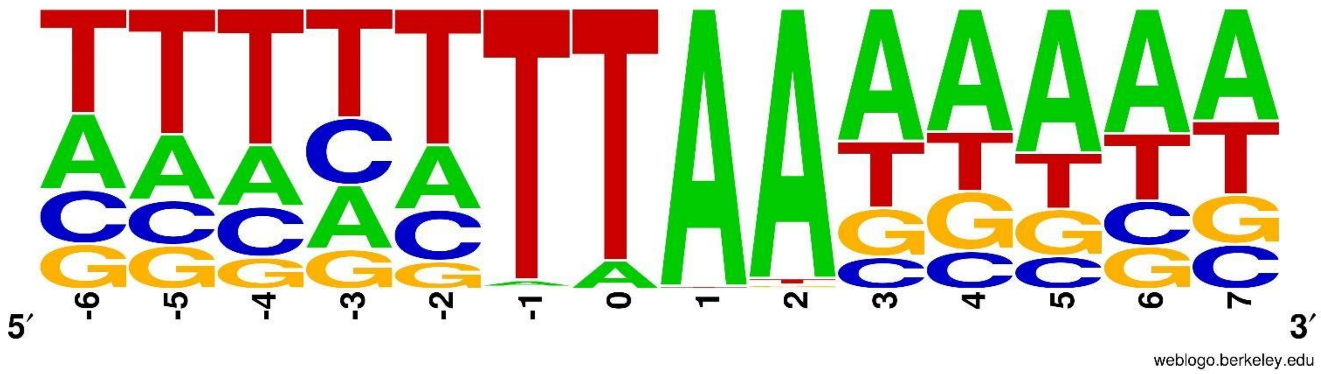


Fig. S16 Logo motif analysis of *CaPB* study.
A graphic representation of the frequency of nucleotides surrounding the *PB* transposon insertions in the *C. albicans* genome

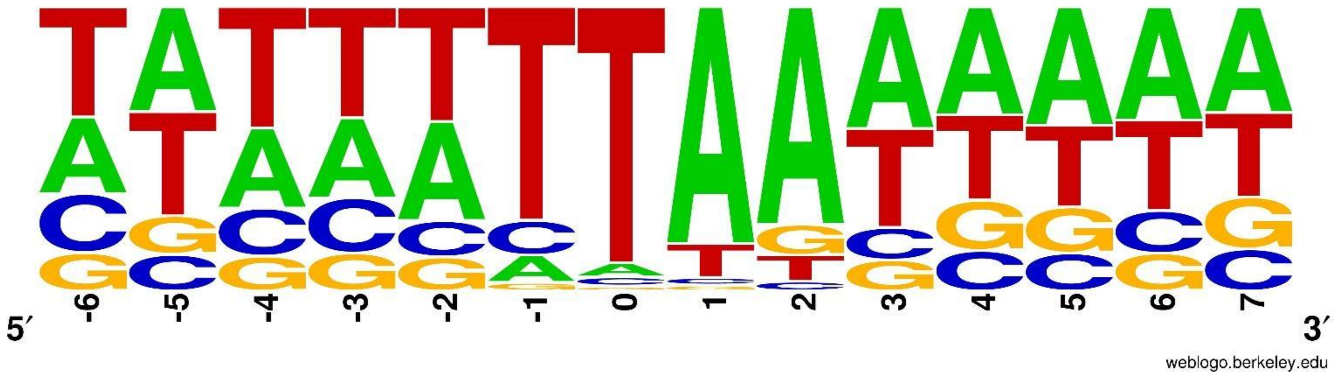

Fig. S17 Comparison of *S. cerevisiae* gene essentiality to Edskes et al., 2018.
Venn diagram of *S. cerevisiae* essential genes, as inferred by three different transposon studies: *ScAc/Ds*, *ScHermes*, and *ScHermes* from Edskes et al., 2018.

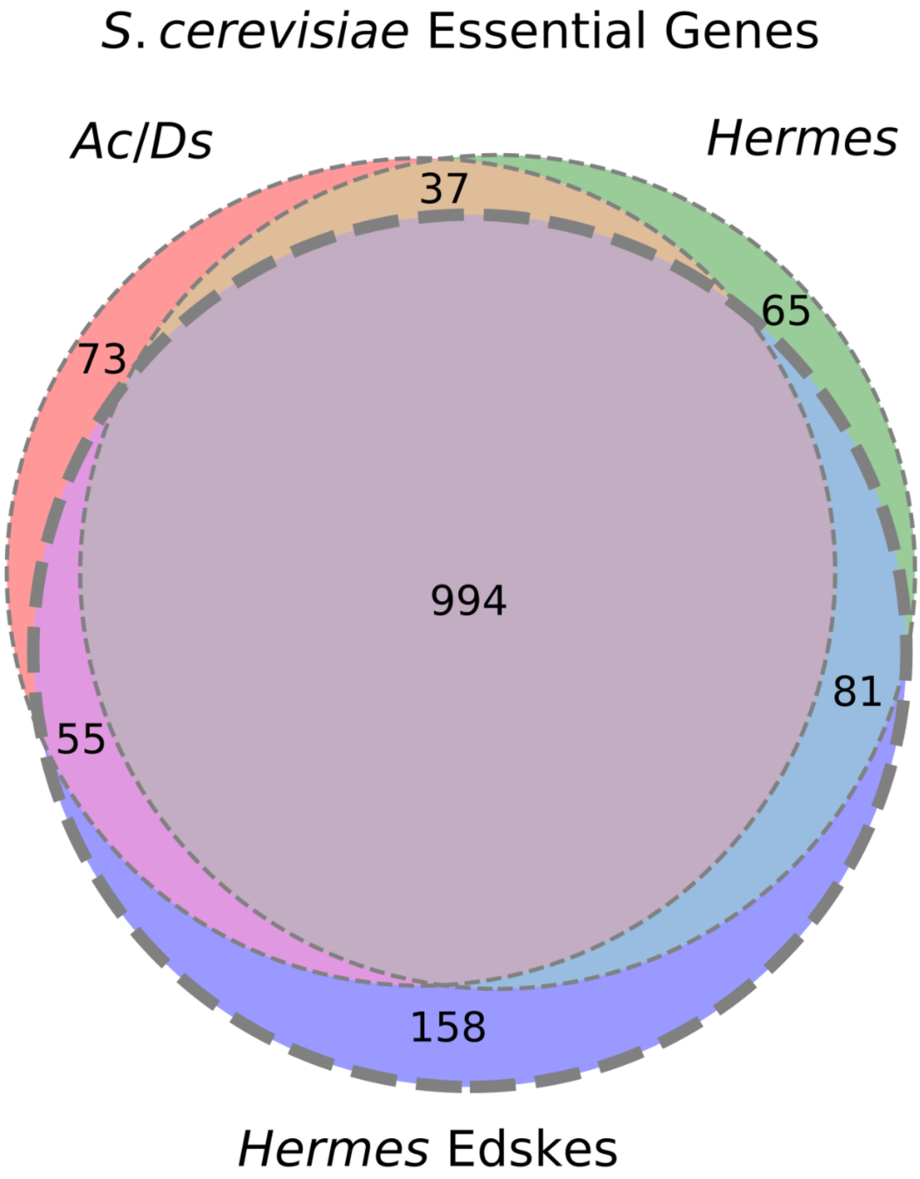
